# Genetic control of the transcriptional response to active tuberculosis disease and treatment

**DOI:** 10.1101/2025.04.11.648383

**Authors:** John F. O’Grady, Alexander S. Leonard, Houcheng Li, Lingzhao Fang, Hubert Pausch, Isobel C. Gormley, Stephen. V. Gordon, David E. MacHugh

## Abstract

Understanding the functional impact of genomic sequence variants is critical for evaluating the role of genetic variation in the host response during tuberculosis (TB) disease and anti-TB treatment (ATT). Hitherto, there have been no genome-wide *in vivo* response expression quantitative trait loci (reQTL) studies conducted for active TB and ATT. Here, using longitudinal peripheral blood RNA-seq data from *n* = 48 patients with active TB who underwent ATT, we call sequence variants directly from these transcriptomes and impute them with a multi-ancestry reference panel. Associating our variants with the expression of nearby genes, we characterise thousands of *cis*-eQTL and hundreds of reQTL. We further show significant changes in cell type proportions during ATT through deconvolution of the bulk RNA-seq data and identify the putative cell type specific nature of *cis*-eQTL. Our work sheds light on the immunogenetics of TB disease and treatment, while providing a framework for studies using only RNA-seq data.

## Introduction

*Mycobacterium tuberculosis* is the principal causative agent of human tuberculosis (TB), an infectious disease that caused the deaths of approximately 1.25 million people in 2023 and with more than 10.8 million newly diagnosed cases during the same reporting period^1^. Worldwide, it is estimated that approximately 25% or circa two billion individuals are latently infected with *M. tuberculosis*^2^. Of these latent TB infected individuals, between 5‒10%, will develop active TB disease that will require treatment^3^. Without treatment, the death rate from human TB disease is high, with estimates ranging from 53-86%^4^; however, close to 85% of cases can be cured with treatment^1^. Anti-TB treatment is a protracted process, characterised by the administration of strong antibiotics during two phases: 1) an intensive phase including rifampicin, isoniazid and pyrazinamide in combination with ethambutol for a period of two months, and 2) a continuation phase encompassing four months of rifampicin or isoniazid^5^. Unfortunately, anti-TB treatment is associated with unwanted adverse effects including fever, hyperuricemia and most notably, hepatotoxicity^6^.

Genetic linkage analyses and more recently, genome-wide association studies (GWAS) have been conducted for numerous tuberculosis-related phenotypic traits reviewed by ^7,8^. These have included studies focusing on anti-TB drug-induced liver injury/hepatotoxicity, albeit with very small case numbers^9,10^. However, as is common to many GWAS studies, genome-wide significant single-nucleotide polymorphisms (SNPs) are largely located in non-coding genomic regions and inherited as dense, highly correlated haplotype blocks making identification of the causal variant challenging^11–13^.

Expression quantitative trait loci (eQTL) are genomic polymorphisms associated with variability in mRNA transcript abundance and can be classified arbitrarily as *cis* if proximal to the associated gene (generally ≤ 1Mb) or *trans* if located further away from the associated gene (> 1Mb, including locations on different chromosomes)^14–16^. Expression QTL are functionally important for polygenic traits such as disease resilience because they are often enriched in GWAS signals^17^ and provide a route for assigning regulatory polymorphisms to complex trait variation^18^. In this regard, previous studies have characterised the regulatory architecture of mammalian genomes, primarily focusing on gene expression across tissues in a range of species (e.g., humans, domestic cattle, and pigs)^19–21^, at key developmental stages^22^, and in particular cell types during specific disease contexts^23^.

In the context of TB disease, several studies have integrated genotype and expression data to explore the genomic architecture of the transcriptional response during TB disease; however, these studies have either focused on a modest number of candidate genes^24^, or were conducted *in vitro* using dendritic cells (DCs) or monocyte-derived macrophages (MDMs)^25–28^. To the best of our knowledge, no high-resolution genome-wide eQTL study has been conducted for TB disease or to profile the genomic architecture *in vivo* of the anti-TB treatment transcriptional response. However, the study by Barreiro *et al.* ^25^ is noteworthy, because genomic variants were identified that alter mRNA expression only upon *M. tuberculosis* infection and likely represent functionally relevant polymorphisms involved in the host response during TB disease. These variants are known as response eQTL (reQTL) and represent a bridge between disease/trait-relevant phenotypes and genetic variation^29^.

Most of the consortia-based eQTL studies outlined above were focused on mapping regulatory loci in bulk tissues. Although this is informative, and given the *cis*-genetic correlation of gene expression across tissues is high at approximately 0.7‒0.8^30^, the cross cell-type *cis*-genetic correlation of gene expression is much lower at circa 0.25‒0.75^31^ indicating cell type specific effects. Recent large-scale studies have therefore focused on mapping such *cis*-eQTL at single-cell resolution (sc-eQTL)^32^. While this is costly and computationally challenging, a method to circumvent these obstacles centres on identifying proxy sc-eQTL known as cell type interacting eQTL (ieQTL) through *in-silico* deconvolution of bulk RNA-seq based on a scRNA-seq reference^33^, and fitting an interaction term in the linear model between genotype and inferred cell type proportions^34,35^. Identifying such ieQTL can characterise additional loci important in complex trait variation that are otherwise masked in bulk RNA-seq^34^. However, hitherto, mapping of sc-eQTL or cell type ieQTL has not been performed in the context of active TB disease or anti-TB treatment *in vivo*.

An eQTL study requires both genotype and gene expression information from the same individual; however, for some studies this is often not the case. In this regard, it has been shown that calling variants directly from high-throughput RNA-seq data can be used to address this problem^36,37^. For example, this approach has been recently used to conduct multi-tissue eQTL analyses in cattle and pigs^20,21^. Moreover, various tools that call variants from short-read sequence data have been developed, including the genome analysis tool kit (GATK)^38^ and the DeepVariant tool^39^, with the latter being specifically modified to call variants from RNA-seq data^40^.

A previous study conducted by Tabone et al. ^41^ examined transcriptional differences between a variety of TB phenotypes ranging from healthy controls to TB patients post anti-TB therapy in peripheral blood (PB) derived from the Leicester TB cohort. The aim of this current study was to leverage the RNA-seq data from Tabone and colleagues^41^ and determine if genetic variability exists that influences the inter-individual transcriptional response during active TB disease and during anti-TB treatment *in vivo*. This hypothesis was tested using PB RNA-seq data, derived from matched individuals with active TB who underwent anti-TB therapy^41^, and by performing RNA-seq variant calling and *in silico* deconvolution to map *cis*-eQTL and ieQTL associated with active TB disease, and to characterise reQTL that alter their effect due to anti-TB treatment *in vivo*.

## Results

### Study design overview and data description

A total of 240 PB RNA-seq samples published by Tabone *et al.* ^41^ from *n* = 48 active TB patients longitudinally sampled across five timepoints (a single untreated timepoint (T0) and four timepoints post commencement of anti-TB treatment (T1‒T4)) were downloaded from the Sequence Read Archive (SRA)^42^ (**Figure 1**). **Table S1** illustrates the Sequence Read Run (SRR) IDs for each of the patients across all five timepoints. **Table S2** details the metadata information for each of the *n* = 48 patients including, age, sex, and day of sampling relative to anti-TB therapy commencement. The median day (D) of sampling relative to anti-TB commencement (± SD) was −1D ± 1.5, +7D ± 2.8, +55D ± 4.2, +166D ± 16, and +286D ± 41 for T0‒T4, respectively (**Figure S1A**; T**able S2**). Based on the self-reported ethnicity of participants, there were 22 South Asian, 14 European, 10 East African, one British-Indian, and one South African individual(s) present (**Figure S1B**). Of the 48 individuals, 30 were male and 18 were female. The mean age of study participants was 37 ± 11 years; for males and females separately, the mean age was 39 ± 11 years and 32 ± 10 years, respectively (**Figure S1C**; **Table S2**).

**Figure 1:**
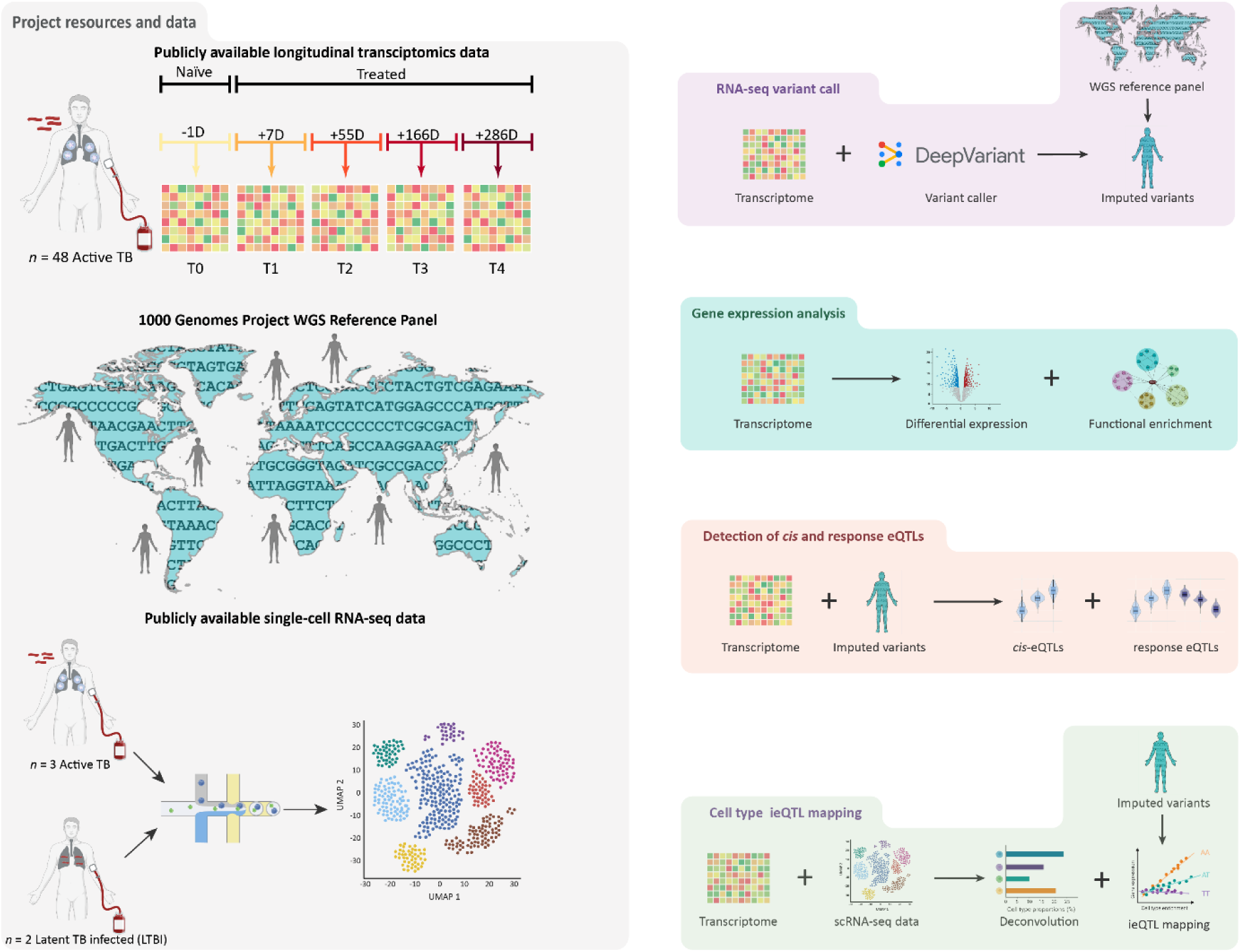
Overview of study design, project resources, and computational analyses. We downloaded publicly available longitudinal peripheral blood (PB) RNA-seq data from ^41^ consisting of *n* = 48 patients with active tuberculosis (TB) who were sampled in a median sense −1D (T0), +7D (T1), +55D (T2), +166D (T3), and +286D (T4), before and after commencing anti-TB treatment. We also obtained whole-genome sequence (WGS) data from 3,202 individuals to act as an imputation reference panel from ^86^ and peripheral blood mononuclear single-cell RNA-seq data from *n* = 3 active TB patients and *n* = 2 latent TB infected (LTBI) patients published by Cai *et al.* ^103^ (grey). The computational genomics analyses included: (1) variant calling with DeepVariant directly from the RNA-seq data and using these for imputation with the 1000 Genomes WGS reference panel (violet); (2) differential expression and functional enrichment analysis between each of the treated timepoints (T1‒T4) and the untreated timepoint (T0) (neon blue); (3) integration of RNA-seq data and imputed variants for *cis*-eQTL and response-eQTL mapping (peach); and (4) integration of bulk RNA-seq, scRNA-seq data and imputed variants for *in silico* deconvolution and cell type interaction eQTL (ieQTL) mapping (mint). Some figure components created with a Biorender.com license.

Peripheral blood RNA-seq post-QC and filtering yielded a mean number of reads per individual sample library of 27,659,544 ± 3,408,460 available for alignment. Of these, 16,872,495 ± 3,323,385 reads mapped uniquely to the human reference genome; 6,967,655 ± 3,189,931 reads mapped to multiple loci (1 < *N* ≤ 10); 173,901 ± 50,990 reads mapped to too many loci (*N >* 10); 3,622,640 ± 1,218510 reads could not be mapped as they were too short; and 22,852 ± 7,672 reads could not be mapped to any genomic locus (**Table S3**). Following quantification of gene expression levels, none of the libraries exhibited an abnormal distribution of counts (**Figure S2A**) and principal component analysis (PCA) based on the top 1,500 most variable genes following variance stabilising transformation of the raw counts in DESeq2 demonstrated that no sample, both within and across time points appeared to be an outlier. However, we observed that principal component 2 (PC2) separated individuals based on their reported sex at timepoints T0‒T2 whereas PC1 separated individuals based on their reported sex at timepoints T3 and T4, respectively (**Figure S2B**). Evaluating the expression of the *RPS4Y1* and *XIST* genes across all *n* = 240 individuals/timepoints demonstrated that samples were assigned the correct sex ID (**Figure S2C**).

### Variant calling from RNA-seq data and imputation

Using DeepVariant, for each individual at each timepoint, we called variants directly from the RNA-seq data and merged timepoint specific samples together using GLnexus. Using this approach, we characterised 1,523,539; 1,602,815; 1,783,843; 1,675,180; and 1,709,479 raw variants for the T0‒T4 groups, respectively. We observed high genotype concordance between paired samples at matched genomic loci, with a mean genotype concordance of 0.9899 ± 0.0125, confirming there were no sample swaps (**Figure S3**; **Table S4**).

To maximise sequence coverage for variant calling, we merged patient-specific BAM files for each of the 48 individuals across timepoints (**Table S5**). The short-read sequence data was generated using RNA sequencing; therefore, the depth at coding sequences (CDS) was high, with about 25% of protein-coding loci covered at a depth of approximately 50× (**Figure S4A**), whereas the coverage across the genome was relatively poor with roughly 5% of the genome covered by approximately two reads (**Figure S4B**). Using the merged BAM files, we called 4,034,779 raw variants across all 48 patients. Of these variants, a total of 791,555, 767,033, 764,644 and 763,894 were present in benchmarked regions of the Genome In A Bottle (GIAB) samples: HG001, HG005, HG006, and HG007, respectively^43^. On average, across all four samples, DeepVariant identified the correct reference and alternative allele pairs for 97.898% of called variants (**Figure S5A**). Removing putative triallelic sites increased the mean percentage of correctly called reference and alternative allele pairs to 99.392% (**Figure S5B**), which demonstrated that DeepVariant can produce highly accurate raw variant calls from PB RNA-seq data.

Following imputation and filtering of variants, a total of 1,506,948 genome-wide variants remained. A principal component analysis (PCA) generated form 22,746 pruned genome-wide variants demonstrated that PC1 and PC2 accounted for 9.33% and 7.47% of the variation observed across the top 20 genotype PCs, respectively, with PC1 separating individuals based on their self-reported African ancestry and PC2 capturing European-Asian differences (**Figure 2A**). We observed that our ability to successfully call and impute variants is largely restricted to expressed coding sequence (CDS) regions as imputation performance decreased substantially in regions of the genome not captured in the RNA-seq alignment (e.g., gene deserts or regions void of expression in these samples) (**Figure 2B**), which is consistent with previous studies^37^.

**Figure 2:**
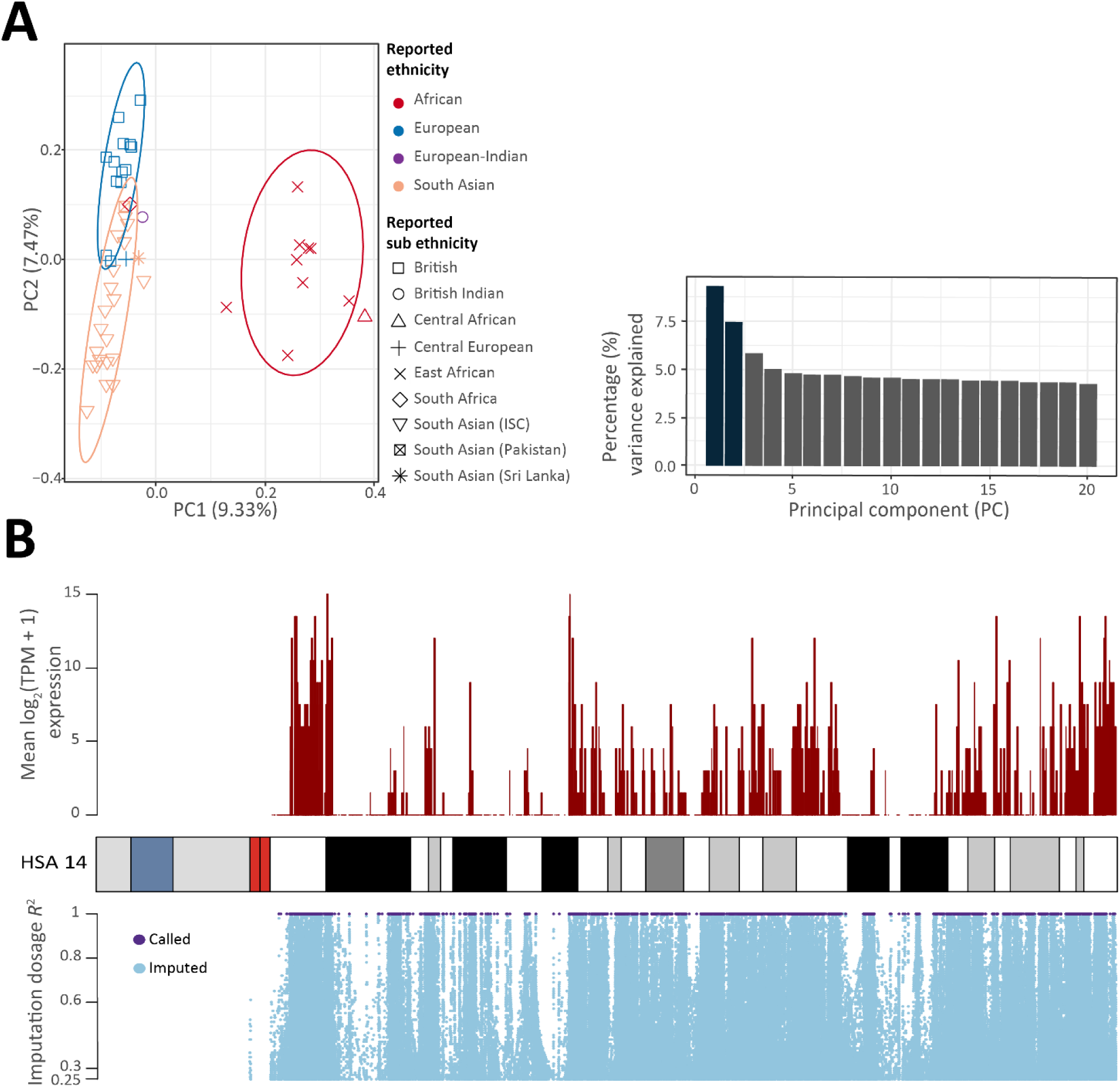
Variant calling from RNA-seq data reveals global geographic structure but high-quality imputation restricted to coding regions. (**A**) principal component analysis (PCA) plot of principal components 1 and 2 (PC1 and PC2) derived from 22,746 pruned imputed variants for the *n* = 48 individuals in this study. The data points are coloured based on the self-reported primary ethnicity of the participants and the shapes represent self-reported sub-ethnicity. A bar plot of the relative variance contributions in the top 20 PCs is also shown with PC1 and PC2 accounting for 9.33% and 7.47% of the variation in the top 20 PCs, respectively. (**B**) A karyoplot of human chromosome 14 (HSA14) showing the relationship between log_2_ (TPM + 1) normalised gene expression from the pooled RNA-seq samples across all *n* = 48 individuals and imputation performance for 267,985 HSA14 variants (Dosage *R*^2^ ≥ 0.25) originally called by DeepVariant (purple) or imputed with Beagle (steel blue).

Taken together, based on these results, we were confident that 1) all samples were labelled correctly, 2) DeepVariant successfully called CDS variants with the correct reference and alternative allele, 3) the genotype data successfully differentiated individuals based on their self-reported ethnicity, and 4) imputation performance in non-coding and non-expressed regions of the genome was suboptimal.

### Differential expression analysis highlights profound transcriptional changes that occur during anti-TB treatment

The original study by Tabone *et al.* ^41^ compared transcriptional perturbations between treated individuals and healthy controls. Here, using a paired design, we tested a related but distinct hypothesis: do significant transcriptional changes occur in PB following anti-TB treatment compared to the same individuals with active TB? We observed that substantial changes occur in the transcriptional profiles of PB during anti-TB treatment with 154 genes significantly differentially expressed (DE) (*P*_adj._ < 0.05, |LFC| > 0.2) at T1 versus T0 (**Figure 3A**). Of these, 77 genes exhibited increased expression and 77 exhibited decreased expression in the T1 group. The number of significant DE genes increased during anti-TB treatment up to a maximum of 1,367 and 1,847 genes that exhibited increased and decreased expression, respectively, in the T3 group versus the T0 group. However, in the comparison between T4 and T0, the number of significant DE genes reduced to 1,212 and 1,592 genes exhibiting increased and decreased expression, respectively, in the T4 group (**Figure 3A**; **Table S6**).

**Figure 3:**
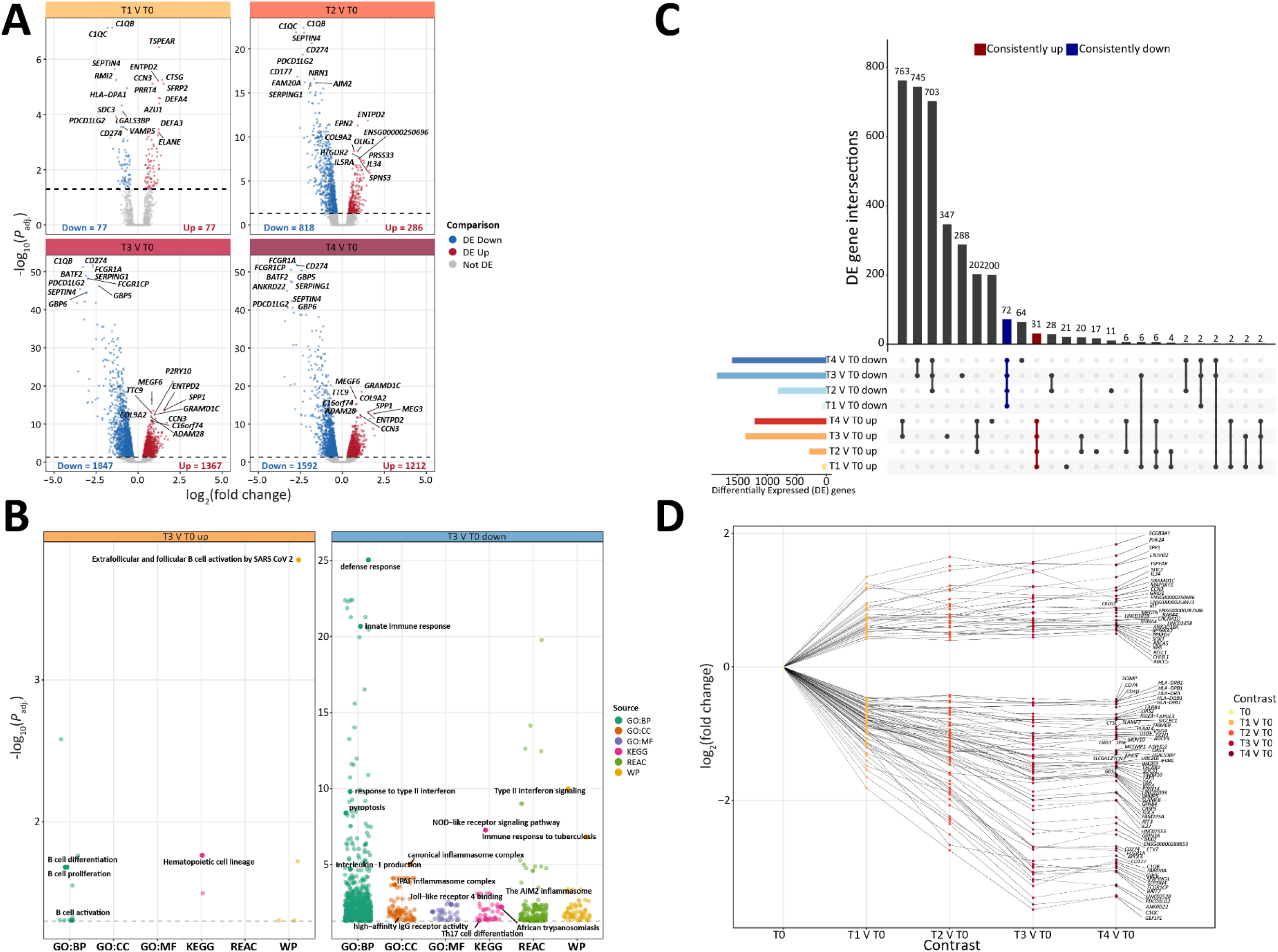
Paired differential expression analysis highlights substantial perturbation of the transcriptome implicating immunologically relevant pathways. (**A**) Volcano plot for each of the differential expression contrasts conducted. The *y*-axis indicates the -log_10_(*P*_adj._) value and the *x*-axis shows the log_2_ fold change (LFC) for each gene between treated conditions (T1‒T4) compared to the untreated condition (T0). Genes significantly increased in expression (*P*_adj._ < 0.05 and LFC > 0.2) are coloured red whereas genes significantly decreased in expression (*P*_adj._ < 0.05 and LFC < 0.2) are coloured blue. Non-significant genes (*P*_adj._ > 0.05) are coloured grey. (**B**) Jitter plot of significantly overrepresented pathways from g:Profiler for the top 300 differentially expressed increased (orange) and decreased (blue) genes from the analysis of T3 versus T0. The *y*-axis shows the -log_10_(*P*_adj._), the *x*-axis and the data point colours illustrates the respective databases from which significant pathways/ontologies originate: GO:BP (Gene Ontology Biological Processes), GO:CC (Gene Ontology Cellular Compartment), GO:MF (Gene Ontology Molecular Function), KEGG (Kyoto Encyclopaedia of Genes and Genomes), REAC (Reactome), and WP (WikiPathways). (**C**) Upset plot showing the overlap of significantly differentially expressed increased (yellow to red) and decreased (light blue to blue) genes across all contrasts. A total of 72 consistently decreased and 31 consistently increased genes are highlighted in dark blue and red, respectively. (**D**) Time-series line plot showing the LFC relative to T0 displayed on the *y*-axis at each contrast displayed on the *x*-axis for 103 consistently significant DE genes (*P*_adj._ < 0.05, |LFC| ≥ 0.2).

In all groups, gene-set enrichment analysis (GSEA) revealed significantly more impacted pathways (SIPs) resulting from genes with increased expression (3025) than from genes with decreased expression (53) (**Figure 3B**; **Table S7**). For genes with decreased expression in the treated groups, the SIPs included the *Immunoglobulin complex* GO:CC ontology (*P*_adj._ = 1.17 × 10^-77^) in the T2 vs. T0 set of genes; the *Immune response to tuberculosis* WikiPathways term (*P*_adj._ = 1.32 × 10^-8^) in the T3 vs. T0 gene set; and the *AIM2 inflammasome complex assembly* GO:BP ontology (*P*_adj._ = 1.23 × 10^-6^) in the T4 vs. T0 gene set. For genes with increased expression, the SIPs included the *LTC4-CYSLTR mediated IL4 production* Reactome term (*P*_adj._ *=* 2.15 × 10^-2^) in the T2 vs. T0 gene set; the *B cell proliferation* GO:BP ontology (*P*_adj._ *=* 2.09 × 10^-2^) in the T3 vs. T0 gene set; and the *B cell receptor signaling pathway* KEGG term (*P*_adj._ = 3.07 × 10^-2^) in the T4 vs. T0 gene set. The complete results for each GSEA are detailed in **Table S7.**

In total, across all contrasts, we identified 7,246 significantly DE genes of which 2,942 exhibited increased expression and 4,334 exhibited decreased expression in the treated groups (T1‒ T4) compared to the naïve untreated group (T0). **Figure 3C** illustrates the overlap of significant DE genes detected within each contrast separated by their direction of change relative to T0. We observed a total of 103 DE genes that were consistently DE in the same direction across all contrasts; 72 were consistently decreased and 31 were consistently increased in the treated groups versus the untreated group (**Figure 3C**, **Figure 3D**). Of these consistently DE genes, 15 genes (*FAM20A*, *CD274*, *CASP5*, *TIFA*, *FCGR1A*, *C1QC*, *SDC3*, *C1QB*, *P2RY14*, *GRAMD1C*, *GBP6*, *PDCD1LG2*, *GRIN3A*, *GBP1P1* and *FCGR1CP*) overlapped with the 27 TREAT-TB gene signature developed by Tabone *et al.* ^41^.

The number of overlapping genes observed was significantly more than what would be expected by chance (*P* < 2.2 × 10^-16^; Fisher’s exact test), supporting the hypothesis that the expressed transcripts of these genes may serve as useful biomarkers for profiling the PB transcriptional response during anti-TB therapy.

### Mapping *cis*-expression QTL reveals the genomic architecture of the transcriptional response during TB disease and anti-TB treatment

Using TensorQTL, we mapped variants proximal to each gene to identify *cis*-eQTL. As *cis*-eQTL generally are arbitrarily defined as being within a ±1 Mb interval relative to the associated gene, our imputed variants are largely restricted to coding regions (**Figure 2B**), and we surmised that this may have implications for selecting an optimum window size for mapping *cis*-eQTL. We therefore selected various *cis* window sizes around the transcriptional start site (TSS) ranging from 0.01 to 1 Mb and mapped corresponding *cis* variants in each group accounting for the top three genotype PCs, age, days since anti-TB treatment commencement, and between six and nine transcriptomic PCs derived from PCA4QTL to account for hidden confounders (**Figure S6**; **Tables S8-S12**).

Assessing the number of unique *cis*-eGenes (i.e., a gene classified as having a significant variant (*P*_adj._ < 0.1) in *cis*, a window size of ± 50 kb away from the TSS was determined to be optimal for the data set **(Figure S7**). In total for this window size, we identified 725, 700, 767, 718, and 748 *cis*-eGenes in the T0, T1, T2, T3, and T4 groups, respectively (*P*_adj._ < 0.1) (**Figure 4A**; **Table S13‒S17**). Of these *cis-*eGenes, between 26.8‒33.7% were timepoint-specific, and 208 were shared across all five groups, which accounted for between 27.1‒29.7% of all *cis*-eGenes detected in each group. (**Figure 4A**). We characterised 6,888, 6,551, 7,828, 6,985, and 7,213 significant *cis*-eVariants with a nominal *P*-value below the genome-wide significance threshold in the T0, T1, T2, T3, and T4 groups, respectively (**Figure S8**). Pleiotropic effects were evident because of these, between 4.0‒7.6% were significantly associated with more than one *cis*-eGene.

**Figure 4:**
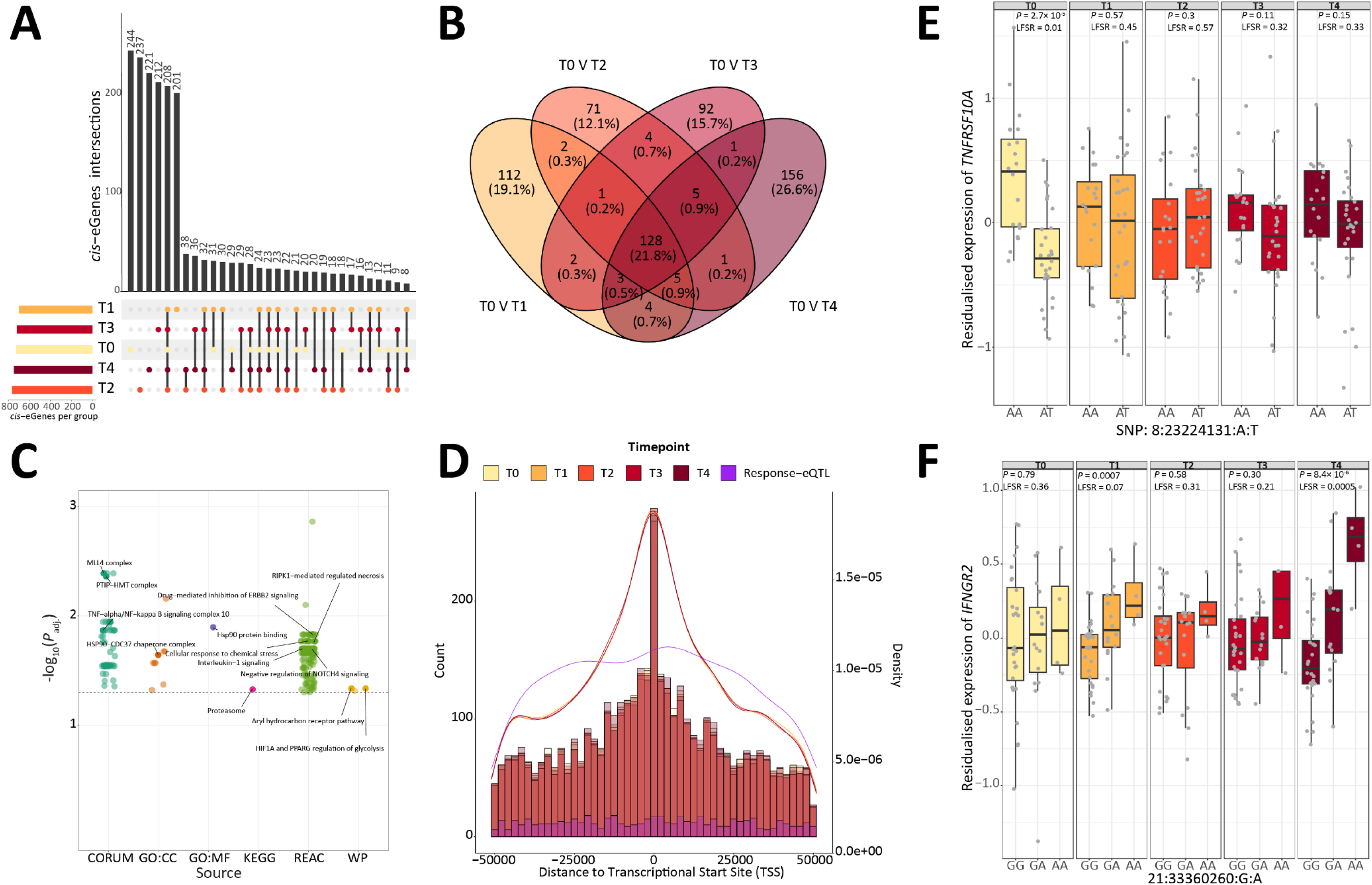
Mapping of *cis* and response-eQTL illustrates genetic control of the transcriptional response to anti-TB treatment in peripheral blood. (**A**) Upset plot showing the overlap of significant *cis*-eGenes (*P*_adj._ < 0.1) among the five timepoints (T0‒T4). The coloured bar plots show the number of *cis*-eGenes detected at each timepoint. (**B**) Venn diagram illustrating the overlap of significant response-eQTL (reQTL) determined from each of the four treatment timepoints (T1‒T4) versus the untreated group (T0). An reQTL was determined as being significant in at least one group (mashr local false sign rate (LFSR) < 0.05) and if the absolute magnitude of fold difference between the two groups in each contrast was ≥ 2. (**C**) Jitter plot of significantly enriched pathways from g:Profiler for all response-eGenes identified. The *y*-axis shows the -log_10_(*P*_adj._) and the *x*-axis and data point colours indicate the pathway/gene ontology databases. (**D**) Distance of all reQTL (purple) and significant *cis*-eQTL (LFSR < 0.05) to the transcriptional start site (TSS) of the target gene identified in each group separately. The overlaid histograms show the distribution in each group of eQTL. The curves show the kernel density estimate for each group of eQTL. (**E**) An example of an active-TB specific response *cis*-eQTL. The bar plot shows the residualised expression for *TNFRSF10A* (See Methods) with individuals plotted based on their genotype at the 8:23224131 locus. Raw *P*-values from the association analysis with TensorQTL and mashr LFSRs are also detailed for each timepoint. (**F**) An example of a treatment-induced response *cis*-eQTL. The bar plot shows the residualised expression for *IFNGR2* with individuals plotted based on their genotype at the 21:33360260 locus. Raw *P*-values from the association analysis with TensorQTL and mashr LFSRs are also detailed for each timepoint.

The power in our study experimental design lies in its longitudinal nature; therefore, we combined information across groups to maximise the statistical power for detecting *cis*-eQTL. To do this, we followed the procedure implemented by Lin et al. ^44^ and used mashr^45^ to recompute effect size and standard error estimates of *cis*-eQTL using inherent correlations shared between the summary statistics from each of the groups. Using this approach, we identified 4,061, 4,049, 3,954, 3,982, and 4,137 significant *cis*-eQTL (local false sign rate (LFSR) < 0.05) in the T0, T1, T2, T3, and T4 groups, respectively (**Table S18**). In total, across all five groups, we identified 5,129 significant *cis*-eQTL of which 3,626 (70.7%) were shared across the five groups. The variance of the posterior mean effect sizes estimated using mashr was significantly lower than the variance of the effect size estimates obtained with TensorQTL (two-sided *F*-test; *P* < 2.2 × 10^-16^) across all groups. Similarly, the posterior standard deviation estimates from mashr were significantly lower than the TensorQTL standard error estimates for each group (two-sided paired *t*-test; *P* < 2.2 × 10^-16^) (**Figure S9**).

We next wanted to characterise variants that change their effect due to the commencement of anti-TB treatment (i.e., reQTL). To do this, we performed pairwise comparisons between the naïve untreated group (T0) and each of the treated groups (T1‒T4) and defined an reQTL as one that was significant in at least one group (LFSR < 0.05), and where the magnitude of the absolute fold difference in the posterior mean effect size of the variant was ≥ 2 in a respective comparison. Using this approach, from each of the four comparisons, we characterised 587 unique reQTL; 431 (73.4%) that were timepoint-specific and 156 (26.6%) that were shared across more than one comparison indicating temporal and persistent effects of genetic variation on gene expression due to the anti-TB treatment (**Figure 4B**; **Table S19‒S22**). To assess if any of the genes associated with these reQTL were involved in biologically important pathways, we performed a GSEA using g:Profiler and ordered genes based on their absolute magnitude fold difference relative to T0. In total, we identified 217 SIPs (*P*_adj_ < 0.05) that encompassed, for example, transcriptional regulation, epigenetic reprogramming, the immune response during *M. tuberculosis* infection, and host response to xenobiotic treatment. These SIPs included the *TNF-alpha/NF-kappa B signaling complexes 6*, *7*, *8*, *9* and *10* (*P*_adj._ ≤ 1.54 × 10^-2^); the *PTIP-HMT complex* (*P*_adj._ = 4.3 × 10^-3^) from the CURAM database; the *HSP90-CDC37 chaperone complex* (*P*_adj._ = 2.1 × 10^-2^) from GO:CC; and the *Aryl hydrocarbon receptor* (*P*_adj._ = 4.6 × 10^-2^), *IPAF inflammasome* (*P*_adj._ = 3.0 × 10^-2^), and *Cellular response to chemical stress* (*P*_adj._ = 1.7 × 10^-2^) Reactome pathways (**Figure 4C**). The complete results from this GSEA are detailed in **Table S23**.

Similar to Lin *et al.* ^44^, we observed that significant *cis*-eQTL (LFSR < 0.05) were enriched around the TSS with no particular trend in the 5’ or 3’ direction, whereas the reQTL tended to be more uniformly distributed across the genomic interval window (**Figure 4D**). However, none of the *cis*-eQTL or reQTL, from any of the groups, were normally distributed around the TSS (Shapiro-Wilk normality test; *P* ≤ 7.8 × 10^-11^ for all QTL sets). Our classification of 587 reQTL allowed us to characterise 137 active TB-specific reQTL that were significant (LFSR < 0.05) only in the T0 group and 438 treatment-induced reQTL that were significant in at least one of the treated groups but not T0. There were no significant reQTL present in all five groups. The active TB-specific response-eGenes included *TNFRSF10A* (**Figure 4E**), *GPX4*, and *NAALADL1*. Response eGenes associated with anti-TB treatment commencement included *RAB27A* at T1, *FBXO3* at T2, *PSMD3* at T3, and *IFNGR2* at T4 (**Figure 4F**).

### Single-cell RNA-seq analysis and computational deconvolution of bulk RNA-seq data

To computationally deconvolve our bulk RNA-seq samples and assess changes in cell type proportions during anti-TB treatment, we first downloaded scRNA-seq data from peripheral blood mononuclear cells (PBMCs) obtained from *n =* 2 latent TB infection (LTBI) patients and *n* = 3 active TB cases. Following filtering of low-quality cells, removal of ambient RNA, and discarding putative doublets, a total of 44,009 cells (6,894‒10,378 cells per sample) remained that were clustered into 15 major annotated cell subtypes (**Figure 5A**). These cells were annotated using a comprehensive procedure involving automatic reference-based annotation with SingleR, marker-assisted annotation using SCINA, and marker-based annotation using published information on cell-type gene markers^46,47^ (**Table S24-25**). **Table S26** details the annotation for each of the 44,009 cells based on the annotation procedure and the final annotation illustrated in **Figure 5A**. From these clusters, we derived a signature matrix of 760 genes to use as reference for the deconvolution of the bulk RNA-seq data (**Table S27)**.

**Figure 5:**
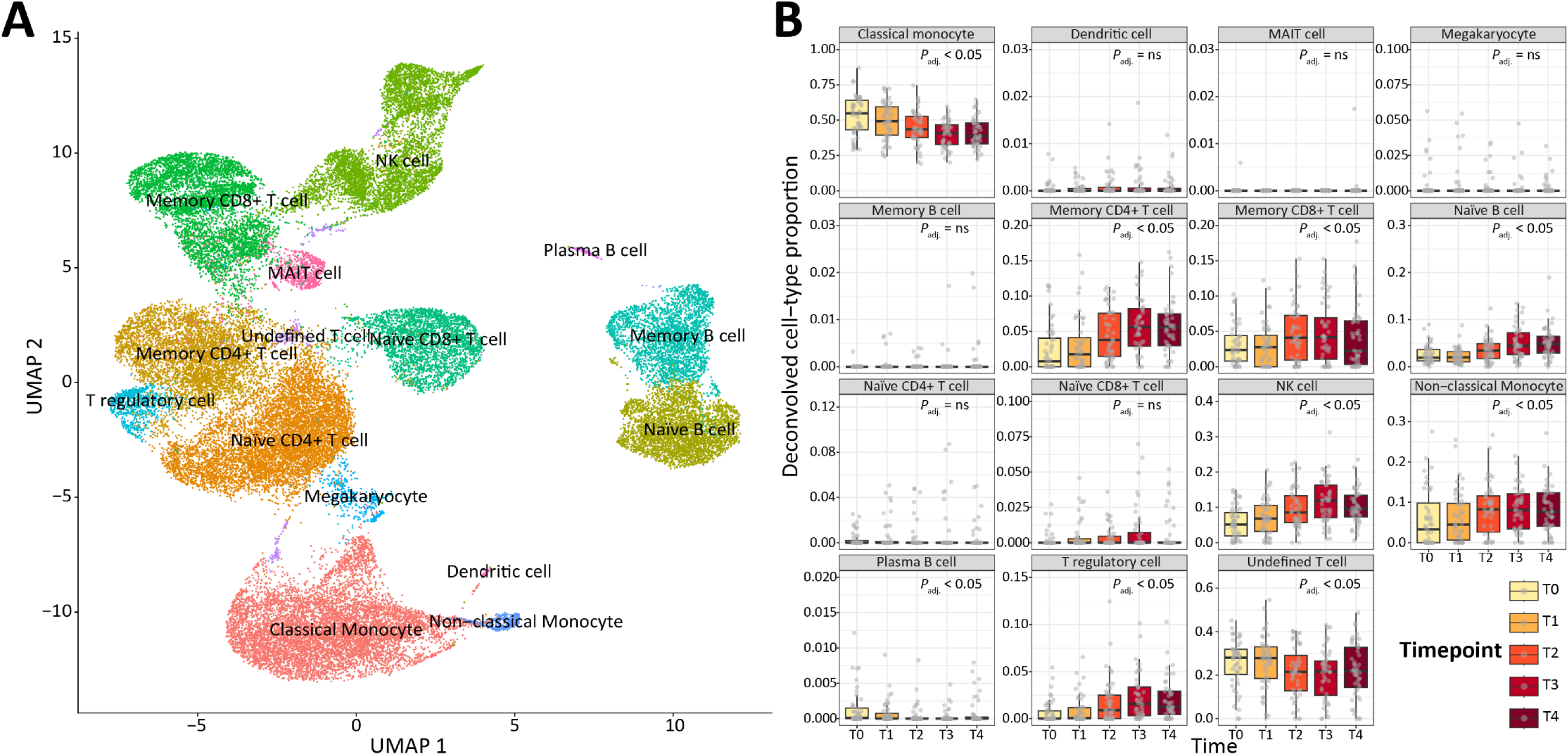
Digital flow cytometry analysis using a scRNA-seq reference shows profound changes in cell type abundance due to anti-TB treatment. (**A**) Uniform manifold approximation and projection (UMAP) of 44,009 cells derived from PBMC scRNA-seq data from *n* = 2 LTBI individuals and *n* = 3 active TB patients. Cells are clustered and coloured into 15 major cell types that are denoted on each cluster. (**B**) Deconvolution results from CIBERSORT for all *n* = 240 bulk tissue samples based on the scRNA-seq reference matrix derived from (**A**) and consisting of 15 major cell types. Significant difference in cell type abundance was inferred based on an ANOVA *F*-test implemented in limma, where *P*_adj._ value < 0.05 indicate significant changes over time and “ns” indicates non-significance. Boxplots are coloured based on sampling timepoint. Horizontal lines inside the boxplots show the medians. Box bounds show the lower quartile (Q1, the 25^th^ percentile) and the upper quartile (Q3, the 75^th^ percentile). Whiskers are minima (Q1 – 1.5 × IQR) and maxima (Q3 + 1.5 × IQR) where IQR is the interquartile range (Q3‒Q1). NK, natural killer; MAIT, mucosal-associated invariant T-cell; and DC, dendritic cell.

Prior to deconvolving the PB RNA-seq data, we performed a benchmarking assessment on pseudo-bulk RNA-seq data for four deconvolution algorithms: Non-negative least squares (NNLS), MuSic, Bisque, and CIBERSORT with various parameter modifications totalling 12 different analyses (See Methods; **Supplementary Note 1**). From this analysis, based on the average Pearson correlation (*ρ*) and average root mean square error (RMSE) estimates across all 15 cell types, CIBERSORT, when specifying the raw scRNA-seq signature matrix and the CPM-normalised pseudo-bulk RNA-seq data, was determined to be the best performing algorithm (*ρ* = 0.998, RMSE = 0.003) (**Supplementary Note 1**).

Based on the results of the deconvolution benchmarking assessment, we deconvolved our CPM-normalised bulk RNA-seq data using CIBERSORT, specifying the signature matrix derived from the scRNA-seq data as the reference. The results of the deconvolution for each of the *n* = 240 bulk RNA-seq samples are presented in **Figure 5B** and **Table S30**. We observed significant changes (ANOVA *F*-test; *P*_adj._ < 0.05) in cell type proportions during anti-TB treatment. For example, natural killer (NK) cells significantly increased over time (*P*_adj._ = 4.92 × 10^-13^), whereas classical monocytes significantly decreased over time (*P*_adj. =_ 7.98 × 10^-12^). Memory CD4^+^ T cells significantly increased over time (*P*_adj._= 4.42 × 10^-13^), as did naïve B cells (*P*_adj._ = 7.92 × 10^-13^) and T regulatory cells (*P*_adj._ = 1.39 × 10^-8^).

Other cell types that displayed significant changes included plasma B cells (*P*_adj._ = 4.49 × 10^-3^), memory CD8^+^ T cells (*P*_adj._ = 1.47 × 10^-2^) and undefined T cells (*P*_adj._ = 2.23 × 10^-2^). The complete results from the PB cell type deconvolution analysis are provided in **Table S31**.

### Cell-type interaction eQTL (ieQTL) mapping

To interrogate the putative cell type specificity of *cis*-eQTL, and to characterise interaction *cis*-eQTL (ieQTL), we leveraged the deconvolved computational abundance estimates of well-characterised and common (median proportion ≥ 0.01 across all timepoints) cell types from CIBERSORT (classical monocytes, non-classical monocytes, memory CD8^+^ T cells, naïve B cells, and NK cells). Fitting an interaction term between genotype and inverse normally transformed cell type abundance estimates in the linear model using TensorQTL accounting for the same covariates used in the standard *cis*-eQTL analysis, we identified no significant interaction terms after multiple testing correction (*P*_adj._ > 0.05). We therefore employed a similar approach to increasing our power in the mapping of *cis*-eQTL in bulk tissue and used mashr to recompute effect size and standard error estimates of this interaction term across timepoints for each cell type separately. We defined a threshold of LFSR < 0.01 as evidence for a significant interaction effect between genotype and cell type abundance.

From this analysis, across all timepoints, we identified a total of 185 significant ieQTL (LFSR < 0.01) in classical monocyte cells. Similarly, across all timepoints, 220, 186, 220, and 110 ieQTL were observed in memory CD8^+^ T cells, naïve B cells, NK cells, and non-classical monocytes, respectively (**Figure 6A**). In total, across all cell types and timepoints, we identified significant interaction terms for 921 variant-gene-cell pairs (**Figure 6A**; **Table S32)**. We observed that ieQTL had a more uniform distribution around the *cis* window, similar to reQTL (**Figure 4D**) across all cell types (**Figure 6B**). Characterisation of significant ieQTL allowed us to identify a total of 115 active TB-specific ieQTL (**Table S32)** across all six cell types tested. For example, we observed a significant classical monocyte-specific ieQTL for *NOD2* (variant = 16:50652646:A:C, *P_g_*_×*c*_ =6.4 × 10^-5^, LFSR = 0.0071) that was only significant at T0 (**Figure 6C**). Other examples of active TB-specific ieQTL included ieQTL at *NLRC4* in classical monocytes (variant = 2:32236868:A:AAAAG, *P*_GxC_ = 0.0003, LFSR = 0.007), *NLRC3* in NK cells (variant = 16:3560641:T:C, *P_g_*_×*c*_ = 0.0002, LFSR = 0.007), and *IFITM2* in memory CD8^+^ T cells (variant = 11:296203:A:G, *P_g_*_×*c*_ = 0.0003, LFSR = 0.003). We also identified treatment-induced ieQTL effects that were not significant in active TB (T0), including a significant ieQTL identified for *ATG4B* in non-classical monocytes early in anti-TB treatment at T1 (variant = 2:241678377:A:G, *P_g_*_×*c*_ = 0.013, LFSR = 0.008) (**Figure 6D**). Other treatment-induced interaction *cis*-Genes included *TNFAIP3*, *ABCA7*, and *S100A6*. The complete results of the ieQTL analysis are detailed **Table S32**. Taken together, our results suggest that cell type-specific *cis-*eQTL are a feature of active TB disease and during anti-TB treatment.

**Figure 6:**
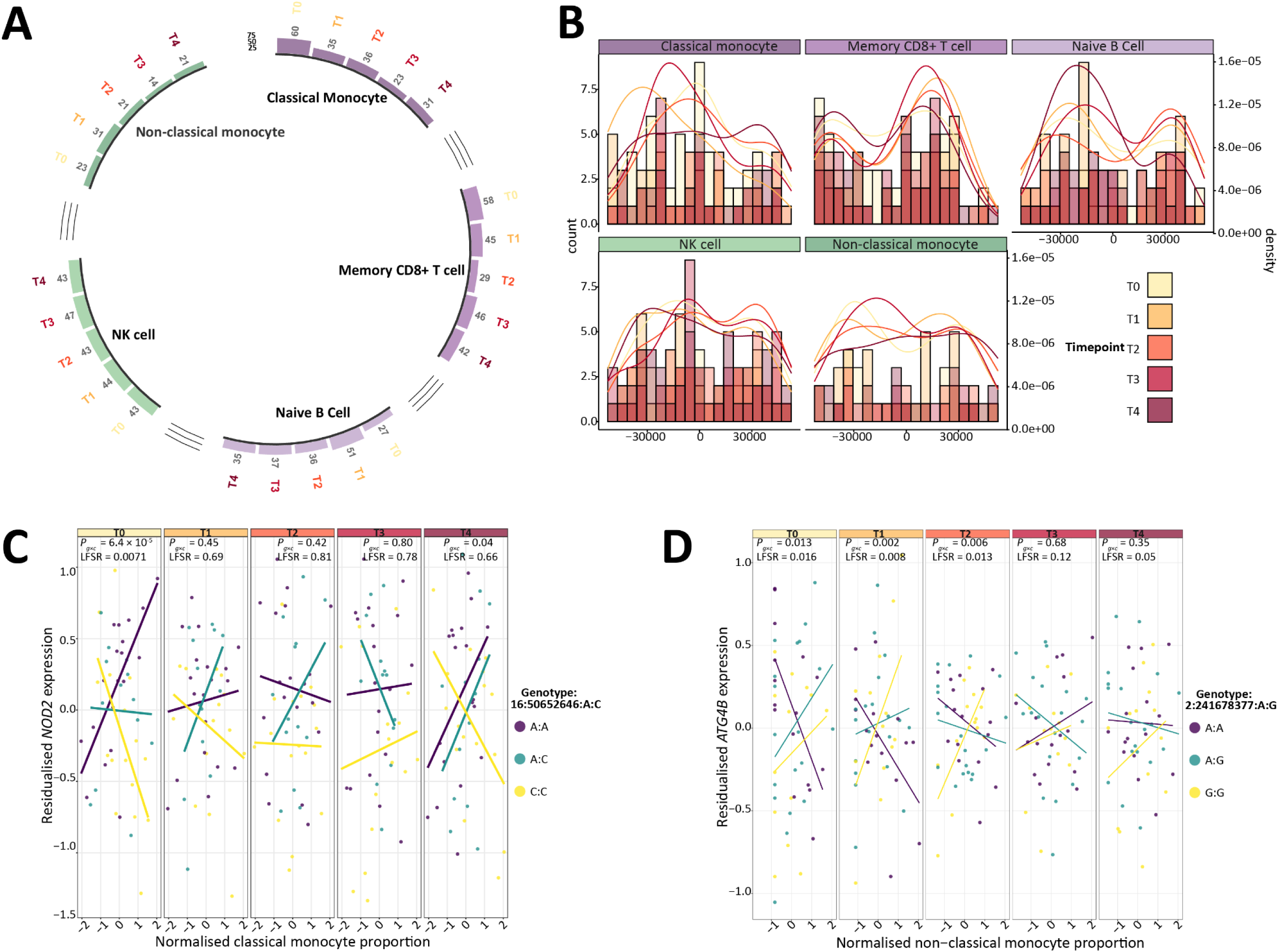
Unearthing the cell type specificity of *cis*-eQTL during active TB and anti-TB treatment through interaction eQTL mapping. (**A**) Circular bar plot showing the number of statistically significant (local false sign rate (LFSR) < 0.01) interaction-eQTL (ieQTL) identified in each cell type (classical monocytes, memory CD8^+^ T cells, naïve B cells, and NK cells) across each of the five timepoints (T0‒T4) separately. (**B**) Histogram organised into 5 kb bins showing distance of the significant ieQTL (LFSR < 0.01) to the transcriptional start site (TSS) of the target gene that were identified in each cell type. The overlaid histograms show the distribution for each timepoint. The curves show the kernel density estimate for each group of eQTL with scales represented on the second *y*-axis. (**C**) Example of an active TB classical monocyte-specific ieQTL identified for *NOD2* with the SNP 16:50652646:A:C. Scatter plot shows the residualised expression values for *NOD2* on the *y*-axis and the inverse normally transformed classical monocyte proportion on the *x*-axis with data points coloured based on corresponding genotype. Solid lines indicate the line of best fit for each genotype derived from the *geom_smooth* function in ggplot2. *P_g_*_×*c*_-values for the interaction term from TensorQTL are also shown as are the LFSR values obtained from mashr. (**D**) Example of a treatment-induced ieQTL identified in non-classical monocytes for the gene *ATG4*B with the SNP 2:241678377A:G in the T1 group. Scatter plot shows the residualised expression values for *NOD2* on the *y*-axis and the inverse normally transformed classical monocyte proportion on the *x*-axis with data points coloured based on corresponding genotype. Solid lines indicate the line of best fit for each genotype derived from the *geom_smooth* function in ggplot2. *P_g_*_×*c*_-values for the interaction term from TensorQTL are also shown, as are the LFSR values obtained from mashr.

## Discussion

Here, we have performed the first *in vivo* high-resolution genome-wide *cis*-eQTL, reQTL, digital flow cytometry, and cell type *cis*-ieQTL mapping analysis in the context of active TB disease and anti-TB treatment using publicly available RNA-seq and scRNA-seq data. Our work illustrates the profound transcriptional and cell type abundance changes that occur during anti-TB treatment and sheds light on how genomic variation influences this response, while also providing a framework for other studies that wish to exploit PB transcriptomics data to its full potential.

### Variant calling from high-resolution transcriptomics data

An abundance of public PB-derived transcriptomics data now exists. For example, in case of human TB, this includes the Berry cohort^48^, the Adolescent Cohort Study (ACS) and the Grand Challenges 6-74 Cohort Study (GC6-74) analysed by Zak et al. ^49^, and the Leicester cohort used for the present study and previously for studying the transcriptional landscape during exposure to *M. tuberculosis*, active TB disease or LTBI, and during or post-anti-TB treatment^41,50^. These data sets have been used primarily for identifying transcriptional biosignatures. Here, however, we show that calling variants directly from these RNA-seq data for the purposes of molecular QTL mapping is a powerful approach to decipher the genomic architecture underpinning the shared and divergent transcriptional responses within each of these phenotypic contexts.

We characterised 1,506,948 high-quality imputed and common variants through pooling of longitudinal RNA-seq data for individual TB patients across timepoints and showed that LD pruning and analyses of these variants recovered known population genetic structure^51^ (**Figure 2A**). However, our ability to impute variants outside of coding regions is significantly curtailed, likely because the imputation algorithm requires an even distribution of variants across the genome, which enables identification of long shared haplotypes in the reference set that can be imputed in the target set with high confidence^52^. In this study, because we are limited to RNA-seq data, our genotyped variants are largely restricted to, and clustered within expressed coding regions that may not conform to default assumptions underpinning genome-wide imputation. We showed that this can impact a *cis*-eQTL analysis, primarily in terms of selecting an optimum *cis* window size (**Figure S6**). Future work to assess imputation strategies that capture extragenic polymorphisms using variants called from RNA-seq data will be important in this regard.

### Expression QTL mapping reveals the genetic architecture of the transcriptional response to anti-TB treatment

Our mapping of reQTL identified hundreds of genomic variants that change their effect due to anti-TB treatment (**Figure 4B**) and we showed that the genes putatively regulated by these reQTL are involved in key immune and drug response pathways such as the *IL-1 signalling pathway*, *cellular response to chemical stress ontology*, and the *aryl hydrocarbon receptor* Wikipathway (**Figure 4C**). The latter pathway is noteworthy, specifically in the context of anti-TB treatment, because in zebrafish (*Danio rerio*) infected *in vivo* with *Mycobacterium marinum* and treated with rifabutin, inhibition of the aryl hydrocarbon receptor pathway with a small molecule inhibitor significantly enhanced bacterial killing compared to controls^53^. Therefore, targeting this pathway in humans, particularly those with variants associated with increased expression of the genes involved, may be a valuable strategy for host-directed therapy and for maximising the effectiveness of current anti-TB therapeutics and development of personalised treatment regimes. Interestingly, similar to Lin *et al.* ^44^, we observed that reQTL tended to be uniformly distributed around the TSS with no particular enrichment at any locus, which is in contrast to standard *cis*-eQTL mapping where these eQTL showed clear enrichment around the TSS (**Figure 4D**). These results suggest that functionally relevant polymorphisms implicated in TB disease and anti-TB treatment response, at least in PB, may be located relatively far from the TSS. This is consistent with previous observations from analysis of DNase I hypersensitive sites, which show a large proportion of trait-associated polymorphisms from human GWAS studies reside in distal gene regulatory elements^54^.

### Computational deconvolution of bulk RNA-seq data is a robust approach for evaluating changes in cell type proportions during anti-TB treatment

Our deconvolution analysis highlighted significant changes in cell type abundance estimates before, and after commencing anti-TB treatment that is consistent with previous flow cytometry analyses (**Figure 5A**). For example, and most notably, the abundance of classical monocytes significantly decreased during anti-TB treatment (*P*_adj._ < 0.05). An increase in monocyte proportions has been documented in patients with pulmonary TB disease^55^ and a high monocyte to lymphocyte ratio (MLR) has been proposed as a potential biomarker for active TB disease^56^. In addition, a meta-analysis conducted by Adane et al. ^57^ demonstrated a significant decrease in the MLR following anti-TB therapy. For the present study, we also observed that NK cells significantly increased their proportion during anti-TB treatment (*P*_adj._ < 0.05), which aligns with previous reports that documented a depletion of NK cells during active TB disease^58^. Similarly, the proportion of memory CD4^+^ T cells significantly increased following anti-TB treatment (*P*_adj._ < 0.05). This observation is in agreement with previous reports that show that the majority (∼ 95%) of antigen-specific effector T cells die following pathogen clearance allowing a pool of memory CD4^+^ T cells to remain (thereby increasing their relative proportion) and provide protection against reinfection^59^. This is further supported by our DE gene results, which showed that many of the genes decreased in expression across all contrasts are involved in proinflammatory immune response pathways (**Figure 3A**; **Figure 3C**) characteristic of active TB, indicating amelioration of the disease^60^. Naïve B cells also significantly increased their proportion (*P*_adj._ < 0.05) following anti-TB treatment, supporting previous observations that highlighted a significant decrease in the frequency of naïve B cells in patients with active TB compared to healthy controls^61^. Again, this is supported by our DE gene results where many of the genes exhibiting increased expression were significantly overrepresented in pathways associated with B cell proliferation and activation **(Figure 3C**; **Table S7**). Taken together, our results demonstrate that *in silico* deconvolution of bulk PB RNA-seq data using CIBERSORT is a powerful and robust approach for evaluating cell-type composition changes during anti-TB treatment.

### Interaction eQTL mapping reveals putative cell-type specificity of *cis*-eQTL

The deconvolved cell type proportions we obtained facilitated identification of 921 variant-gene pairs significantly interacting with the cell-type abundance estimates for the five cell types tested. Of note, we identified a private, active TB specific ieQTL (16:50652646:A:C) in classical monocytes for the nucleotide binding oligomerization domain containing 2 gene (*NOD2*) (**Figure 6C, Table S32**) that was not considered a *cis*-eVariant in the standard *cis*-eQTL mapping. Specifically, we observed that individuals with the alternative allele displayed lower expression of *NOD2* in classical monocytes compared to heterozygotes or individuals homozygous for the alternative allele. The NOD2 protein is a member of the NLR (NOD, leucine-rich repeat (LRR)-containing protein) family of intracellular pattern recognition receptors through its ability to recognise bacterial cell wall components known as muramyl dipeptides^62^. In TB disease, NOD2 plays a central role in recognising *M. tuberculosis* pathogen-associated molecular patterns (PAMPs), and is essential for the activation of innate and adaptive immune responses to mycobacterial challenge through crosstalk with Toll-like receptors (TLRs)^63^. In this regard, numerous studies across diverse populations have suggested that polymorphisms in the *NOD2* gene are associated with susceptibility to TB^64^. Also, *NOD2*^-/-^ mice experimentally infected with *M. tuberculosis* displayed higher bacterial burdens versus wild type mice, had impaired inflammatory responses and reduced concentrations of TNF and IFN-γ cytokines^65^. Considered together, our results suggest that individuals who are homozygous for the alternative allele of the SNP 16:50652646:A:C have reduced expression of *NOD2* in classical monocytes during active TB disease, which may impede their ability to recognise mycobacterial PAMPs upon phagocytosis and contribute to *M. tuberculosis* spread and pathogenesis.

Our results highlighted several treatment-induced ieQTL including the SNP 2:241678377:A:G associated with the expression of the autophagy related 4B cysteine peptidase gene (*ATG4B*) in non-classical monocytes for which the interaction term was significant at T1 (LFSR = 0.007) and that also tended towards significance at T0 (LFSR = 0.014) and T2 (LFSR = 0.011) (**Figure 6D**). The *ATG4B* gene is a member of the ATG4 gene family encoding proteases that enable efficient autophagosome maturation and cellular autophagy, primarily through delipidation of ATG8 family members that attach to a developing phagophore membrane^66,67^. Considered primarily to be a homeostatic process, autophagy principally involves the breakdown of damaged cellular material through encasement in a double layered membrane organelle known as an autophagosome, which is then degraded further through fusion with an acidic lysosome^68^. Notwithstanding the role of autophagy in homeostasis, it has also emerged, over the past two decades, as a key host defence strategy against invading microbes, including tuberculous mycobacteria^69,70^. For example, in murine macrophages infected with *M. tuberculosis*, inhibition of autophagy through depletion of ATG5 resulted in substantial necrosis and increased host susceptibility to *M. tuberculosis* infection^71^. In the context of *ATG4B*, reduction of *ATG4B* expression levels has been shown to lead to changes in autophagy flux^66^ and in murine macrophages infected with the *M. bovis* Bacillus Calmette-Guérin (BCG) strain, *Atg4b* expression and autophagy were shown to be inhibited by a micro-RNA (miR-129-3p) leading to enhanced intracellular survival of *M. bovis* BCG^72^. Collectively, our results indicate that individuals homozygous for the reference allele at the SNP 2:241678377:A:G have reduced expression of *ATG4B* in non-classical monocytes, which may hinder the autophagic capacity of these cells in the early stages of anti-TB treatment thereby stunting the elimination of *M. tuberculosis*.

### Limitations of the study design

There are necessary and inherent limitations associated with our analyses that should be acknowledged. Firstly, given that we have *n* = 48 samples per timepoint, even after using mashr to improve power, our ability to identify *cis*-eQTL is limited to those with large effects. Secondly, our *cis*-eQTL analysis is restricted to ±50 kb from the TSS. As a result, there may be independent upstream or downstream *cis*-eQTL/reQTL/ieQTL located further than 50 kb from the TSS, which influence the expression of the associated gene that we were unable to identify. Thirdly, while GWAS for anti-TB drug induced liver injury have been conducted, these were generated using relatively small sample sizes and summary statistics are not publicly available. As further GWAS are conducted for this trait and made publicly available, our results will form an important resource to colocalise^73^ putative *cis*-eQTL or reQTL or to perform an integrative transcriptome-wide association study (TWAS)^74^ to characterise the functional genomic mechanisms underpinning this trait. Fourthly, *trans*-eQTL have been shown to contribute to a substantial proportion of gene expression heritability, more so than *cis*-eQTL^75^ and represent an integral component of the omnigenic model of complex trait variability^76^. The relatively small sample size in the present study meant that we were unable to characterise distal genomic variants associated with variability in mRNA transcript abundance. However, exploiting the full set of individuals in the Berry^48^ and Leicester cohorts^41,50^ using our approach may facilitate the identification of *trans*-eQTL within an active TB disease context. Lastly, our deconvolution analysis focused on broad high-level cell types (e.g., classical monocytes and NK cells). As more active TB-specific and anti-TB treatment-specific scRNA-seq data becomes available coupled with advancements in cell type annotation, *in silico* deconvolution can be performed at higher resolution, which would enable characterisation of significant changes in the fine-grained proportions of cell types due to anti-TB xenobiotic treatment.

### Conclusion

Our high-resolution *cis*-eQTL, reQTL, and cell type interaction eQTL analysis has identified factors systematically linking host genomic variation, cell type abundance, and the inter-individual transcriptomic response during active TB disease and anti-TB treatment. Our results form an important resource for researchers investigating how genetic variability impacts the inter-individual response to *M. tuberculosis* infection and anti-TB treatment while also providing a computational framework for other studies that are limited to RNA-seq data alone. Further integrative genomics work, linking GWAS and molecular quantitative trait loci data together is required to fully elucidate the genomic architecture of active TB disease and the responses to anti-TB therapeutics.

## Methods

### RNA-seq sample download and preprocessing

A total of 240 high-throughput paired-end (2 × 100 bp) peripheral blood (PB) RNA-seq fastq files generated by Tabone *et al.* ^41^ derived from a total of *n* = 48 individuals across five timepoints were downloaded from the Sequence Read Archive^42^ (GEO accession number: GSE157657) using fastq-dump (v. 2.10.5) in addition to the *--split-files* parameter. The quality of the RNA-seq reads was assessed using FastQC (v. 0.11.9)^77^ and MultiQC (v. 1.9)^78^. Trimmomatic (v. 0.38)^79^ was used to sequentially remove Illumina adapters, filter raw reads < 36 bp in length and remove trailing bases with a phred-scale quality score < 30. Filtered RNA-seq reads were aligned to the human reference genome (GRCh38_full_analysis_set_plus_decoy_hla) using STAR (v. 2.7.10b)^80^. The reference genome index was generated using the *genomeGenerate* command with the gencode.v43.annotation.gtf annotation file specified in addition to the --*sjdbOverhang* 99 parameter. Read counts for each gene were quantified using featureCounts (v. 2.0.3)^81^ by specifying the *-p -B - C -s2* flags to indicate that paired-end reads were being analysed, to only count pairs with both ends aligned, to exclude chimeric fragments and to denote that reversely stranded RNA-seq reads were being analysed. We performed expression quantification for each timepoint separately resulting in a total of five count matrices.

### RNA-seq variant calling and genotype concordance

Single-nucleotide polymorphisms (SNPs) and indels were called from the aligned reads for each timepoint separately using the DeepVariant (v. 1.5)^39^ RNA-seq variant calling pipeline^40^, including the RNA-seq model from DeepVariant and by specifying the *--split_skip_reads* parameter. Individual sample genomic VCF files derived from DeepVariant were merged and harmonized using GLnexus^82^ generating timepoint-specific VCF files. Across each timepoint separately, we retained variants with a minor allele frequency (MAF) > 0.1, which did not deviate significantly from Hardy-Weinberg Equilibrium (HWE) (*P* < 1 × 10^-6^), and that possessed a genotype call rate > 95%. The genotype concordance of samples across timepoints at matched genomic loci was compared to ensure that no sample swaps occurred. Genotype concordance was defined as the proportion of genotypes (0/0, 0/1, 1/1) at identical loci that matched between any two samples across two timepoints. We compared the genotype concordance for all timepoint pairwise comparisons (20 in total).

To maximise genome coverage and depth, and to generate a final set of variants for each of the *n* = 48 samples, we used SAMtools (v 1.15.1)^83^ to merge individual specific aligned bam files across timepoints together. The two exceptions to this method included samples for patient 3 and patient 157, which possessed a moderately lower genotype concordance at timepoint 1 and timepoint 0, respectively, which was hypothesised to be due to reduced RNA quality. Therefore, for these two samples, we excluded these specific timepoints from the merging process. We then reimplemented the DeepVariant/GLnexus RNA-seq variant calling pipeline on the merged bam files. Read coverage at coding sequences was assessed using mosdepth (v. 0.3.3)^84^. To ensure variants (reference and alternative alleles) were being called correctly, we compared locus-specific reference and alternative allele pairs called in our dataset that were present in benchmarked regions across four Genome in a Bottle (GIAB) data sets (HG001, HG005, HG006 and HG007; https://ftp-trace.ncbi.nlm.nih.gov/giab/ftp/release/) (v. 4.2.1)^43^.

### Phasing and imputation

Prior to phasing and imputation, we excluded rare variants (MAF < 0.03), which deviated significantly from HWE (*P* < 1 × 10^-5^) or that displayed a poor Phred quality score (QUAL < 20) and that were missing in more than 80% of samples. Phasing was performed on each chromosome separately using SHAPEIT (v 5.1)^85^ with individual chromosome recombination maps specified (https://github.com/odelaneau/shapeit4/blob/master/maps/genetic_maps.b38.tar.gz) in addition to a phased reference panel derived from Byrska-Bishop et al. ^86^ composed of 3,202 WGS (30× sequencing depth) individuals. Imputation was performed on each chromosome separately using Beagle (v. 5.4)^87^ with default settings implemented and specifying the same reference panel as used for the phasing procedure. We visualised the relationships between gene expression, variant calling, and imputation performance using the karyoplotR package^88^. Briefly we first plotted the log_2_ transcripts per million (TPM) + 1^89,90^ expression values of genes on chromosome 14 (HSA14) and visualised the dosage *R*^2^ (DR2) values of directly called and imputed variants on the same chromosome. Following imputation, we retained variants with a DR2 value > 0.6 and that had a MAF > 0.05.

Population genetic structure was assessed using the *--pca* parameter from PLINK (v. 1.9)^91^. The *--keep-allele-order*, and *--indep-pairwise 50 10 0.1* flags were used to remove variants in high linkage disequilibrium (LD). The orthogonal eigenvectors of principal components (PCs) 1 and 2 were plotted and data points were coloured according to the self-reported ancestry for each individual.

### Differential expression and functional enrichment analyses

A paired DEA was conducted between the naïve, untreated group (T0) and each treated group (T1, T2, T3, and T4) separately using DESeq2 (v.1.40.2)^92^ and a design matrix incorporating the patient ID as a covariate. Genes with raw expression counts ≥ 6 in at least 20% of samples were retained prior to the DEA. For each DEA, the null hypothesis tested was that the absolute logarithmic fold change (LFC) between the untreated timepoint and each of the treated timepoints for the expression of a particular gene was greater than 0.2. For these contrasts, raw *P*-values were inferred using the Wald test and corrected using the Benjamini-Hochberg (BH) method^93^ to account for multiple tests. Genes with a BH false discovery rate (FDR) adjusted *P*-value (*P*_adj._) < 0.05 and an absolute LFC > 0.2 were considered significantly differently expressed (DE).

To identify pathways/ontologies enriched by significant DE genes, for each contrast, we took the top 300 genes showing increased and decreased expression based on *P*_adj._ and performed a gene-set enrichment analysis (GSEA) using the g:GOSt function implemented in the g:Profiler2 R package (v. 0.2.3)^94,95^. Of note, for DE genes identified from the T1 versus T0 comparison, we did not perform this analysis as the number of input DE genes was too small (*n* = 153) Following best practice recommendations from Wijesooriya et al. ^96^, the background set of genes consisted of all genes that were tested in the DE analysis. Raw *P*-values were corrected using the BH method and pathways with a *P*_adj._ value < 0.05 were considered significantly overrepresented with DE genes.

### *Cis*-eQTL mapping

For the mapping of *cis*-eQTL, we used the human Genotype Tissue Expression (GTEx) Consortium^19^ eQTL analysis pipeline with some minor modifications. We conducted the *cis*-eQTL analysis on each experimental group (T0, T1, T2, T3, and T4) separately. Raw RNA-seq read counts were normalised using the trimmed mean of the M values (TMM) method^97^ and the expression values for each gene were then inverse normally transformed across samples to ensure the molecular phenotypes followed a normal distribution. Genes with raw expression counts ≥ 6 and a TPM^89,90^ normalised expression count ≥ 0.1 in at least 20% of samples were retained for the eQTL analysis. For each group, we used the PCAForQTL R package (v.0.1.0)^98^ to identify hidden confounders in the normalised and filtered expression matrices. We used the *runElbow* function in PCAForQTL to select the number of latent variables using the elbow method. We then merged these inferred/latent covariates with known covariates (the top three genotype PCs of the imputed genotype data set, age in years, and days since anti-TB treatment commencement) and removed highly correlated known covariates captured well by the inferred covariates (unadjusted *R*^2^ ≥ 0.9) using the PCAForQTL *filterKnownCovariates* function.

For the *cis*-eQTL mapping procedure, we used TensorQTL (v.1.0.8)^99^. To the best of our knowledge, no previous published study has investigated the optimal genomic interval at which to conduct a *cis*-eQTL analysis using imputed variants originally derived from RNA-seq data. Therefore, we conducted the *cis*-eQTL analysis considering distance windows of ±10‒1000 kb from the TSS and selected the window size that resulted in the maximum number of unique *cis*-eGenes (i.e., genes with at least one significant *cis*-eQTL) across all groups for downstream analyses. To identify significant *cis*-eQTL, we invoked the permutation strategy in TensorQTL^100^ to estimate variant-phenotype associated empirical *P-*values with the parameter --mode *cis* to account for multiple variants being tested per molecular phenotype. We then used the Storey and Tibshirani FDR procedure^101^ to correct the *beta* distribution-extrapolated empirical *P-*values to account for multiple phenotypes being tested genome-wide. A gene with at least one significant associated *cis*-eQTL (FDR *P*_adj._ < 0.1) was considered a TensorQTL *cis*-eGene.

To identify significant *cis*-eVariants associated with detected *cis*-eGenes, we followed the procedure implemented by the Pig GTEx Consortium^21^. Briefly, we first obtained nominal *P*-values of association for each variant-gene pair using the parameter --mode *cis_nominal*. We then defined the empirical *P*-value of a gene which was closest to an FDR of 0.1 as the genome wide empirical *P-* value threshold (*pt*). Next, we calculated the gene-level threshold for each gene from the beta distribution by using the *qbeta(pt*, *beta_shape1*, *beta_shape2*) command in R (v. 4.3.2)^102^ with beta_shape1 and beta_shape2 being derived from TensorQTL. Variants with a nominal *P*-value of association below the gene-level threshold were included in the final list of variant-gene pairs and were considered as significant TensorQTL *cis*-eVariants.

### Response eQTL (reQTL) mapping

To identify variants that significantly alter their effect due to anti-TB treatment, we followed the procedure implemented by Lin *et al.* ^44^. Briefly, we used mashr (v. 0.2.79)^45^, which maximises power through jointly analysing genetic effects on gene expression across timepoints to improve effect size and reduce standard error estimates facilitating the identification of both shared and condition specific *cis*-eQTL. Following the mashr eQTL analysis outline pipeline (https://stephenslab.github.io/mashr/articles/eQTL_outline.html), we first extracted a “strong set” of effect size estimates derived from the top eQTL for each gene across timepoints to prevent double counting of eQTL that are in LD and impact the same gene. We then extracted a subset of 200,000 randomly selected variants and considered a variant as a *cis*-eQTL and as having a significant effect on gene expression if the local false sign rate (LFSR) was < 0.05.

We next compared the effect sizes of putative eQTL between T0 and each of the treated timepoints (T1‒T4). We classified an eQTL as a response eQTL if the effect was significant in at least one group (LFSR < 0.05) and if the absolute log_2_ fold change in the posterior effect size was greater than 2^44^.

### Enrichment analysis of reQTL

We used the *g:GOSt* function implemented in the g:Profiler2 R package (v. 0.2.3)^94,95^ to determine the biological processes impacted by response-eGenes. To do this, we selected all response-eGenes and ordered them based on their absolute magnitude fold change relative to T0 and removed duplicate genes. The background set of genes consisted of all genes that were tested in the reQTL analysis. Raw *P*-values were corrected using the BH method and pathways with a *P*_adj._ Value < 0.05 were considered significantly overrepresented with response-eGenes genes.

### Single cell RNA-seq data analysis

#### Alignment and quality control

A total of *n* = 5 peripheral blood mononuclear cell (PBMC) scRNA-seq data sets derived from *n* = 3 individuals with active tuberculosis and *n* = 2 individuals with latent tuberculosis infection (LTBI) were obtained from Cai et al. ^103^ and downloaded from the SRA (accession no. <u>SRP247583</u>; Sequence Read Run (SRR) IDs SRR11038989‒SRR11038993). For each sample, the cleaned data filtered for low-quality reads and unrelated sequences were imported to CellRanger (v. 7.2.0)^104^ and aligned to the human reference genome (hg38, GRCh38) specifying the count command and setting the *–expect-cells* parameter to 10,000. The resulting count matrix files were loaded into R using the *Read10x* function from the Seurat (v 5.0.2) R package^105^. Ambient RNA was first removed in each sample using the *autoEstCont* function in SoupX (v.1.6.2)^106^. To remove possible doublets, we used scDblFinder (v. 1.17.1)^107^ setting the dbr parameter equal to 0.01. Finally, we filtered and removed low-quality cells with < 500 or > 3000 expressed genes or that that had >7% of UMIs mapping to mitochondrial genes.

#### Batch effect correction and dimension reduction

Using the Seurat R package, data were log-transformed using the *NormalizeData* function and the top 2000 variable genes were selected using the *FindVariableFeatures* function setting the selection method to “vst”. To remove batch effects, we first obtained genes that were variable across samples using the *SelectIntegrationFeatures* function and identified anchors using the *FindIntegrationAnchors* function setting the reduction parameter = “cca”. Finally, we used the *IntegrateData* function to merge different samples together. The top 12 eigenvectors derived from the *RunPCA* function of the integrated data were used for downstream uniform manifold approximation and projection (UMAP) visualization and clustering.

#### Cell clustering, marker gene identification, and cell type annotation

The Louvain algorithm was used to cluster cells setting the resolution parameter to 0.4 in the *FindClusters* function resulting in 16 distinct cell clusters. We annotated cells using a comprehensive procedure involving automatic reference-based annotation, marker-based automatic annotation and marker gene-based annotation using marker genes curated from the literature. For the reference-based annotation, we used SingleR (v. 2.2.0)^108^, specifying the Human Primary Cell Atlas (HPCA)^109^, Blueprint/ENCODE (BPEN)^110,111^, Immunological Genome Project (ImmGen)^112^ and Monaco immune (Monaco)^113^ data sets as references from the celldex (v. 1.10.1)^108^ package and extracted both the main and high-resolution (fine) cell-type annotation. For the marker-assisted annotation, we used SCINA^114^ specifying markers derived from PBMCs collated by Diaz-Mejia et al. ^115^ in .gmt format. Lastly, we used marker genes that have been systematically curated^46,47^.

To identify marker genes capable of demarcating each annotated cluster, we used the *FindConservedMarkers* for each cluster, grouping on disease status and only considering positively DE genes. A gene was considered significantly DE if it was expressed in at least 25% of the cells within the cluster, the difference in the fraction of detection to the other clusters was at least 0.25 and the LFC was > 0.75. The one exception to this was for *Undefined T cells* where we relaxed the logfc.threshold parameter to 0.5 and the min.diff.pct parameter to 0.15. We then grouped DE genes together based on their cluster, ordered by LFC and selected up to a maximum of 100 genes per cluster and removed duplicated genes. The resulting signature matrix was used in the downstream deconvolution analysis where specified.

### Benchmarking of bulk RNA-seq deconvolution strategies

To identify a suitable tool to deconvolve the bulk RNA-seq data using the scRNA-seq data from Cai *et al.* ^103^ as a reference, we performed a deconvolution benchmarking assessment of four algorithms: CIBERSORT^33^, MuSiC (v 1.0.0) and non-negative least squares (NNLS) (v 1.0.0)^116^, and Bisque (v. 1.0.5)^117^. We first generated pseudo-bulk RNA-seq data representing *n* = 100 samples derived from the Cai *et al.* ^103^ PBMC scRNA-seq data using the SimBu R package (v. 1.2.0)^118^ setting the *scenario* parameter to mirror_db and the *balance_even_mirror_scenario* parameter to 0.5 to introduce variability. We then visualised the cell type proportions of the 100 pseudo-bulk samples using the *plot_simulation* function. Given that MuSiC and Bisque perform internal normalisation, but CIBERSORT does not (unless quantile normalisation is specified that is recommended for microarray data), we separately TPM- and counts per million (CPM-) normalised the pseudo-bulk RNA-seq samples used for the CIBERSORT deconvolution procedure. Additionally, we also separately TPM-normalised the scRNA-seq data prior to deconvolution benchmarking using CIBERSORT.

In total, across all algorithms and normalisation strategies, we performed 12 benchmarking assessments. This included MuSiC, NNLS, and Bisque with, and without specifying marker genes from the signature matrix, and CIBERSORT with the raw, TPM- or CPM-normalised bulk RNA-seq data and the raw or TPM-normalised scRNA-seq data. Comparing the observed cell type proportions post-deconvolution to the expected cell type proportions, we assessed deconvolution performance using two metrics: 1) the average Pearson correlation (*ρ*) and 2) the average inverse of the root mean squared error (RMSE) across cell types where the RMSE is calculated as:

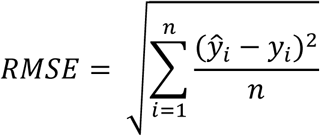

Where *ŷ*_1_, *ŷ*_2_,…, *ŷ*_n_ = are the predicted values and *y*_1,_ *y*_2_,…, *y*_n_ are the true simulated values in the pseudo-bulk samples for a particular cell type.

### Deconvolution of bulk RNA-seq data

To perform the actual deconvolution of the bulk RNA-seq data, we used CIBERSORT^33^ with the CPM-normalised bulk data and the raw counts from the signature matrix generated in this study. We disabled batch correction, quantile normalisation and conducted the analysis in relative mode specifying 100 permutations. Deconvolved cell type proportions were visualised in R using ggplot2 (v. 3.4.4)^119^.

To test for significant differences in cell type proportions throughout treatment, we first normalised the deconvolved cell type proportions using the ascrin method as part of the propeller function derived from the speckle R package^120^. We then used the *lmFit* function in limma (v. 3.56.2)^121^ to fit a linear model accounting for individual ID as a random variable. To test for significant differences across groups, we performed an ANOVA using the *topTable* function and corrected for multiple testing using the BH method. The null hypothesis tested here was that difference in cell type proportions for a particular cell between all the timepoints is equal to 0. Cell types with an FDR *P*_adj._ < 0.05 were considered to significantly change their cell type proportions over time.

### Interaction eQTL (ieQTL) mapping

To identify putative variants that modulate RNA transcript abundance within a particular cell type, we used TensorQTL to fit a linear model that included an interaction term between genotype and cell type proportions that accounted for the same covariates used in the standard *cis*-eQTL mapping procedure. Considering only well-defined cell types with a mean proportion > 0.01 across all timepoints (NK cells, classical monocytes, non-classical monocytes, naïve B cells and memory CD8^+^ T cells), we first normalised the proportions of each cell type across samples using the inverse normal transformation procedure. Within the regression framework, to mitigate against potential regression outlier effects, we restricted ieQTL mapping to variants with MAF ≥ 0.05 in the samples belonging to each of the top and bottom halves of the cell-type proportion distribution, for each timepoint-cell type combination using the *--maf_threshold_interaction* 0.05 option in TensorQTL^34^. To identify genes in each group with at least one significant ieQTL, the top nominal *P-*value for each gene within each cell type analysis was selected and corrected for multiple testing at the gene level using eigenMT^122^. Genome-wide significance was then determined by computing BH FDR on the eigenMT-corrected *P*-values. Following this approach, there were no interaction terms significant after multiple testing correction (*P*_adj._ < 0.05).

To leverage the longitudinal nature of this study and to increase power, for each cell type, we used mashr^45^ to combine information across timepoints and improve effect size of putative ieQTL. To do this we first filtered for common variant-gene pairs tested in the standard analysis using TensorQTL across all timepoints for a particular cell-type. Following the mashr eQTL analysis outline pipeline (https://stephenslab.github.io/mashr/articles/eQTL_outline.html), for each cell type, we selected the lead variant-gene pair across timepoints as the “strong subset” of associations. Next, we selected a random set of 200,000 variant-gene associations to act as the “random set” and invoked a threshold of LFSR < 0.01 to establish if an interaction effect was significant within a particular timepoint.

### Visualisation of eQTL, reQTL, and ieQTL results

To plot results of the eQTL, reQTL, and ieQTL analyses, we extracted the residualised expression levels of genes using the QTLtools (v. 1.3.1)^123^ *correct* function specifying the phenotype .bed file with the parameter *--bed* and the covariates .txt file with the *--cov* parameter. For genes of interest, we plotted the residualised expression values for each individual separated by their corresponding genotype as a boxplot for the eQTL, and reQTL analyses. For the ieQTL analysis, we plotted the residualised expression values and the inverse normally transformed cell type proportions as a scatter plot with data points coloured according to an individual’s corresponding genotype for a particular locus.

## Supplemental Table Legends

**Table S1:** This table reports the patient and SRR IDs for peripheral blood bulk RNA-seq data downloaded from Tabone *et al*., (2021) and used in the analysis, related to Figure 1.

**Table S2:** This table reports the metadata for all *n* = 48 patients, related to Figure 1 and Figure S1.

**Table S3:** This table reports the RNA-seq alignment statistics, related to Figure 1 and Figure S2.

**Table S4:** This table reports the genotype concordance between samples across timepoints, related to Figure 2 and Figure S3.

**Table S5:** This table reports the merged SRR IDs for each sample for the final variant call, related to Figure 2 and Figures S3-5.

**Table S6:** This table reports the differential expression results for each contrast, related to Figure 3.

**Table S7:** This table reports the enrichment results from g:Profiler for the top (most significant) 300 differentially expressed genes exhibiting increased and decreased expression, respectively, related to Figure 3.

**Table S8:** This table reports the covariates used for each sample in the *cis*-eQTL analysis in the T0 cohort, related to Figure 4 and Figure S6-7.

**Table S9:** This table reports the covariates used for each sample in the *cis*-eQTL analysis in the T1 cohort, related to Figure 4 and Figure S6-7.

**Table S10:** This table reports the covariates used for each sample in the *cis*-eQTL analysis in the T2 cohort, related to Figure 4 and Figure S6-7.

**Table S11:** This table reports the covariates used for each sample in the *cis*-eQTL analysis in the T3 cohort, related to Figure 4 and Figure S6-7.

**Table S12:** This table reports the covariates used for each sample in the *cis*-eQTL analysis in the T4 cohort, related to Figure 4 and Figure S6-7.

**Table S13:** This table reports the TensorQTL permutation results for T0, related to Figure 4 and S6-9.

**Table S14:** This table reports the TensorQTL permutation results for T1, related to Figure 4 and S6-9.

**Table S15:** This table reports the TensorQTL permutation results for T2, related to Figure 4 and S6-9.

**Table S16:** This table reports the TensorQTL permutation results for T3, related to Figure 4 and S6-9.

**Table S17:** This table reports the TensorQTL permutation results for T4, related to Figure 4 and S6-9.

**Table S18:** This table reports the significant mashr results, related to Figure 4 and S9

**Table S19:** This table reports the response-eQTL results for the T0 V T1 comparison, related to Figure 4 and S9.

**Table S20:** This table reports the response-eQTL results for the T0 V T2 comparison, related to Figure 4 and S9.

**Table S21:** This table reports the response-eQTL results for the T0 V T3 comparison, related to Figure 4 and S9.

**Table S22:** This table reports the response-eQTL results for the T0 V T4 comparison, related to Figure 4 and S9.

**Table S23:** This table reports the enrichment analysis of response *cis*-eGenes, related to Figure 4.

**Table S24:** This table reports the markers genes used from Karlsson *et al*., 2021 used for annotating cells from the scRNA-seq data, related to Figure 5.

**Table S25:** This table reports the markers genes used from Oelen *et al*., 2022 used for annotating cells from the scRNA-seq data, related to Figure 5.

**Table S26:** This table reports the cell type clustering and annotation for all 44,009 cells, related to Figure 5.

**Table S27:** This table reports the signature matrix of key genes used in the deconvolution analyses, related to Figure 5 and S10-13, and Supplemental Note 1.

**Table S28:** This table reports the pseudobulk data for 100 samples, related to Figure S10-13 and Supplemental Note 1.

**Table S29:** This table reports the deconvolution benchmarking results of different sorftware tools and parameters based on the 100 pseudobulk samples, related to Figure S10-13 and Supplemental Note 1.

**Table S30:** This table reports the deconvolution results of the real bulk RNA-seq data from *n* = 240 samples, related to Figure 5.

**Table S31:** This table reports the Anova *F*-test results for differences in cell type proportions, related to Figure 5.

**Table S32**: This table reports the interaction eQTL results for common cell types, related to Figure 6

## Resource availability

### Lead contact

Requests for further information and resources should be directed to and will be fulfilled by the lead contact, David MacHugh.

### Materials availability

This study did not generate new unique reagents.

### Data and code availability

The *n* = 240 RNA-seq paired-end fastq files derived from the *n* = 48 individuals are available from the Sequence Read Archive (SRA) (GEO accession number: <u>GSE157657</u>) ^41^. The reference genome used for the alignment of the RNA-seq reads can be found at the following link: GRCh38_full_analysis_set_plus_decoy_hla. The DeepVariant variant calling model ^39,40^ is available at the following link: https://github.com/google/deepvariant. The GIAB data sets can be accessed at the following address: https://ftp-trace.ncbi.nlm.nih.gov/ReferenceSamples/giab/release. The PBMC scRNA-seq data from *n* = 3 active TB patients and *n* = 2 LTBI patients used in this study are available on the SRA (accession number <u>SRP247583</u>; SRR IDs SRR11038989‒SRR11038993) ^103^. The filtered and imputed variants for all *n* = 48 samples in addition to the raw *cis*-eQTL, reQTL and ieQTL results will be made publicly available upon publication. The computer code and scripts used in this study are available at the following GitHub link: https://github.com/jfogrady1/Human_TB_reQTL.

## Acknowledgements

J.F.O’G was supported by Research Ireland through the Research Ireland Centre for Research Training in Genomics Data Science (grant no. 18/CRT/6214). This study was also supported by Science Foundation Ireland (SFI) Investigator Programme Awards to D.E.M. and S.V.G. (grant nos. SFI/08/IN.1/B2038 and SFI/15/IA/3154), and University College Dublin (UCD) College of Health and Agricultural Sciences seed funding awarded to J.F.O’G. Note: Since the 1^st^ of August 2024, Science Foundation Ireland (SFI) has been part of Taighde Éireann – Research Ireland (www.researchireland.ie).

The authors would like to thank Joe Keane, John Browne, Thomas Hall, Adnan Khan, and Kseniia Maximova for useful discussions.

## Author contributions

J.F.O’G., A.S.L., H.P., I.C.G, and D.E.M. conceived and designed the study. J.F.O’G., S.V.G., I.C.G. and D.E.M. acquired funding for the study. J.F.O’G. and A.S.L. performed bioinformatics analyses in collaboration with L.F., H.L., and H.P. J.F.O’G. wrote the first draft of the manuscript, and prepared all figures, tables, and supplementary information with input from D.E.M. All authors read and approved the final manuscript.

## Declaration of interests

The authors declare no competing interests.

## Consent for publication

Not applicable

## Notes

### Competing Interest Statement

The authors have declared no competing interest.

