## Supplemental Information for "Genetic control of the transcriptional response to active tuberculosis disease and treatment"

### Supplementary Figures

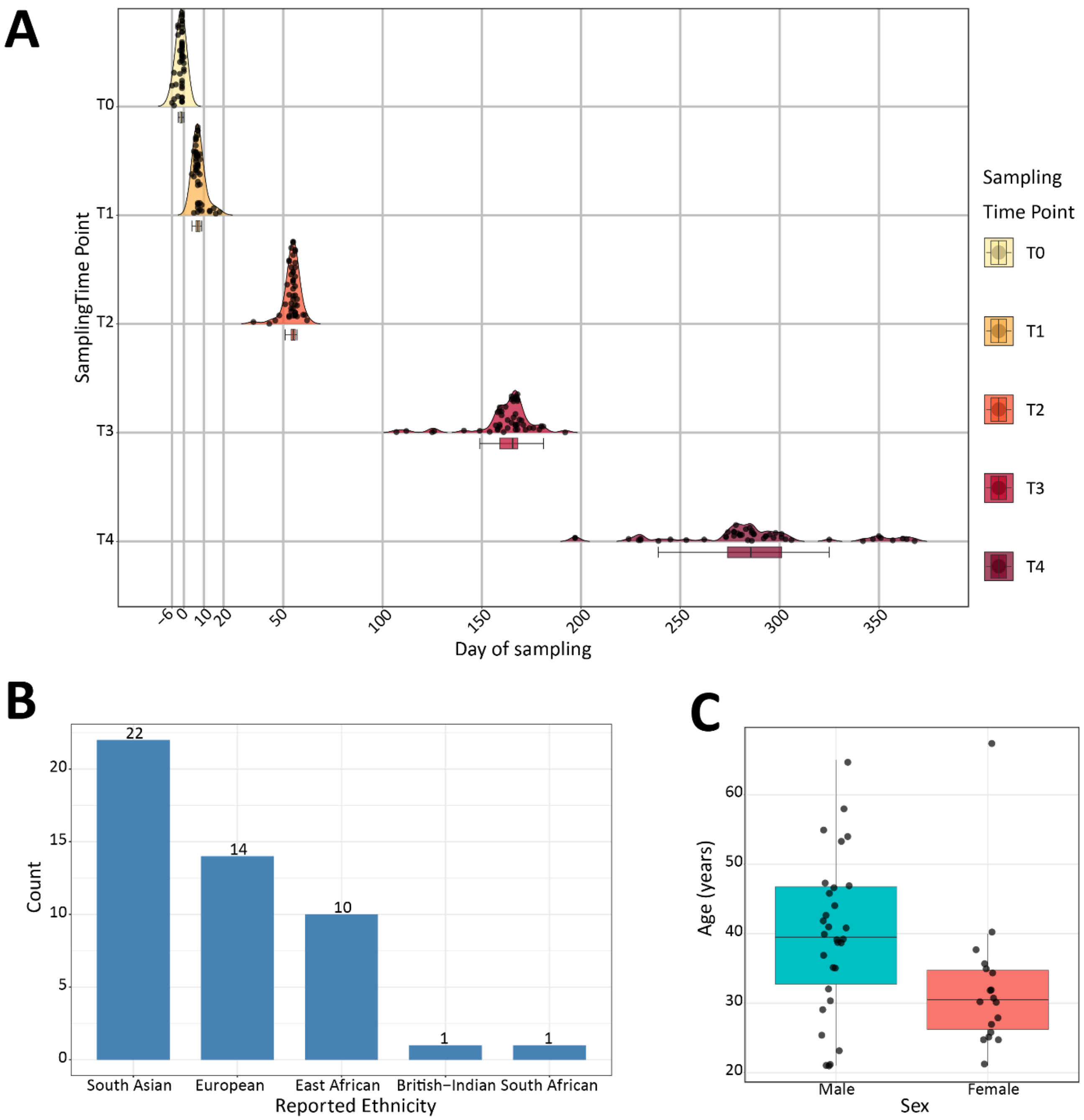

**Figure S1: Sample metadata information, related to Figure 1.**
(**A**) Semi-raincloud plot showing the distribution of day of sampling for all *n* = 48 individuals across all five timepoints analysed (T0–T4). The line in the box plots indicates the median. Whiskers are minima (Q1 − 1.5 × IQR) and maxima (Q3 + 1.5 × IQR), where IQR is the interquartile range (Q3–Q1). (**B**) Bar plot showing the reported ethnicity of all *n* = 48 participants. (**C**) Box plot showing the age distribution of participants plotted by sex. The median of the distributions is indicated by a solid horizontal line within the box plot. Whiskers are minima (Q1 − 1.5 × IQR) and maxima (Q3 + 1.5 × IQR) where IQR is the interquartile range (Q3–Q1).

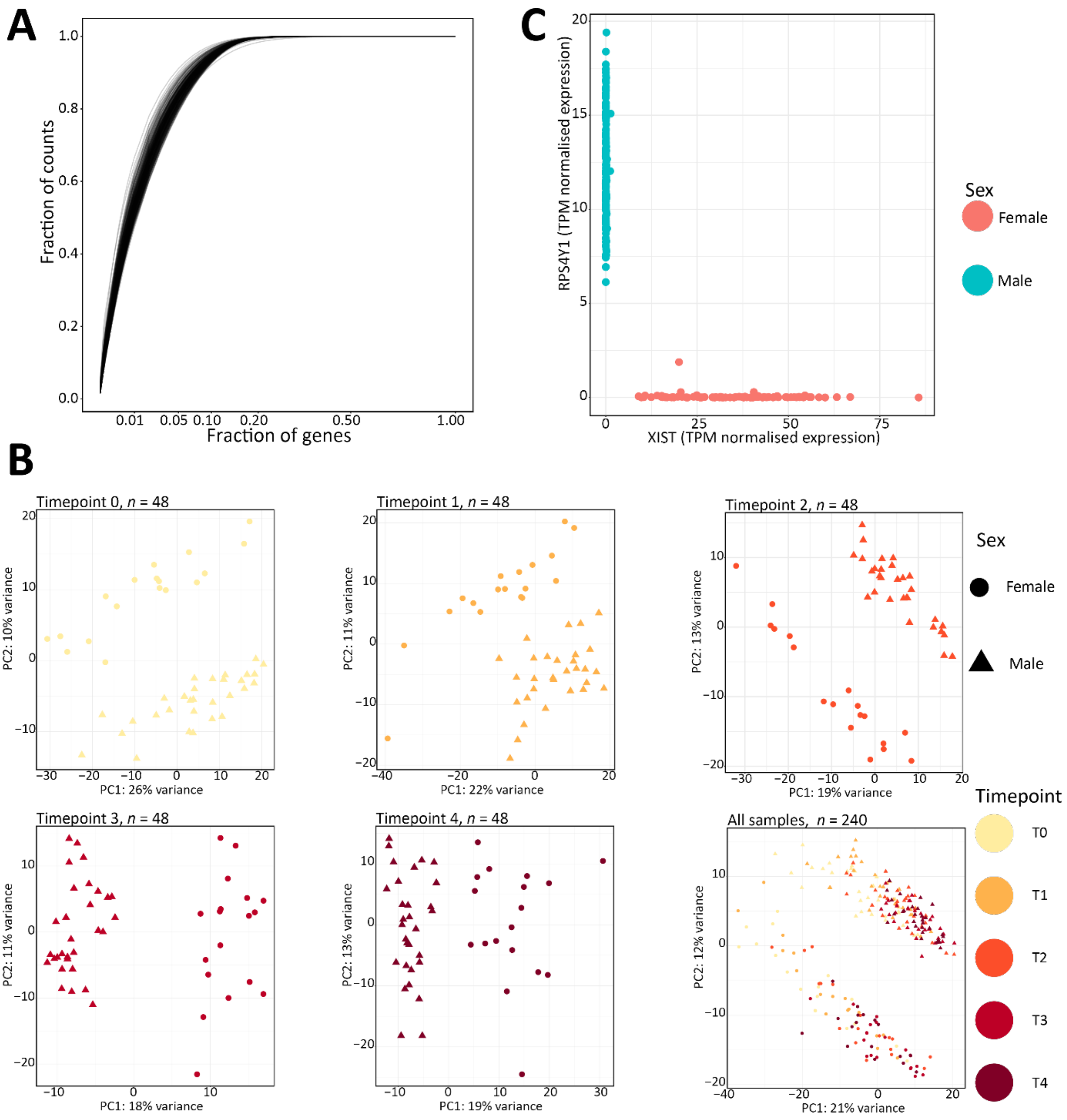

**Figure S2: RNA-seq quality control, related to Figure 1.**

(**A**) Line plot showing the fraction of counts captured by the proportion of genes after expression quantification in each library (*n* = 240). (**B**) Principal component analysis (PCA) of the top 1500 most variable genes as determined from the variance stabilised transformed RNA-seq data in DESeq2[1] for each timepoint and all RNA-seq samples (*n* = 240) together with principal components 1 and 2 (PC1 and PC2) shown in each comparison. Samples are coloured based on their designated cohort. (**C**) Scatter plot showing the relationship between a female-specific gene (*XIST*) and a Y-linked gene (*RPS4Y1*) with individuals coloured based on their reported sex.

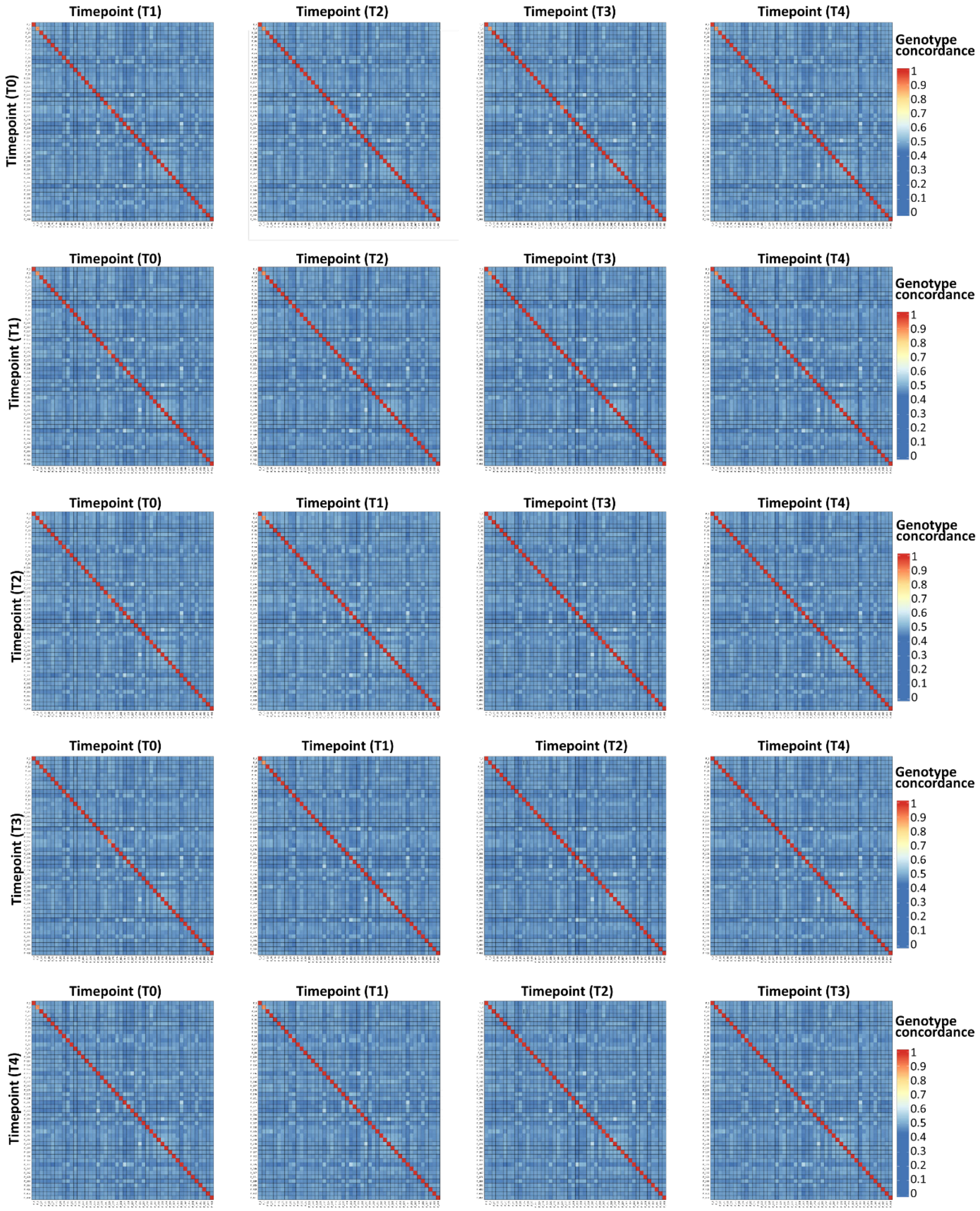

**Figure S3: Sample mismatch assessment, related to Figure 1.**
Genotype concordance using raw variants that were called from all individuals ($n$ = 48) in timepoint 0 (T0), timepoint 1 (T1), timepoint 2 (T2), timepoint 3 (T3), and timepoint 4 (T4) versus all individuals in the other cohorts, respectively. A high genotype concordance indicates a sample match.

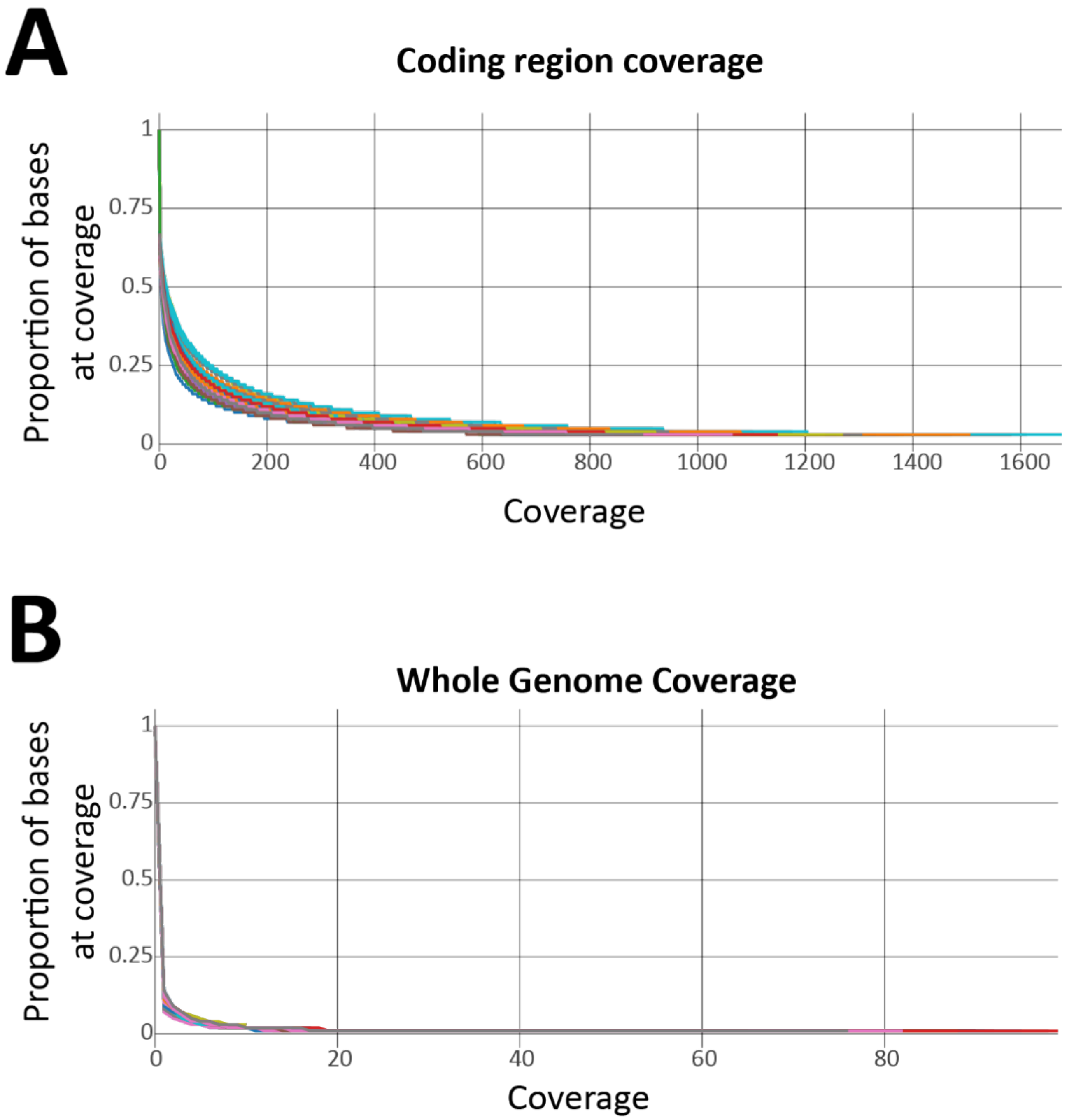

**Figure S4: Variant call coverage assessment, related to Figure 2.**
(**A**) Plot showing the relationship between the proportion of bases covered in coding regions and at what coverage using the merged BAM files for all *n* = 48 individuals. (**B**) Same as (**A**) but displaying coverage across the entire genome.

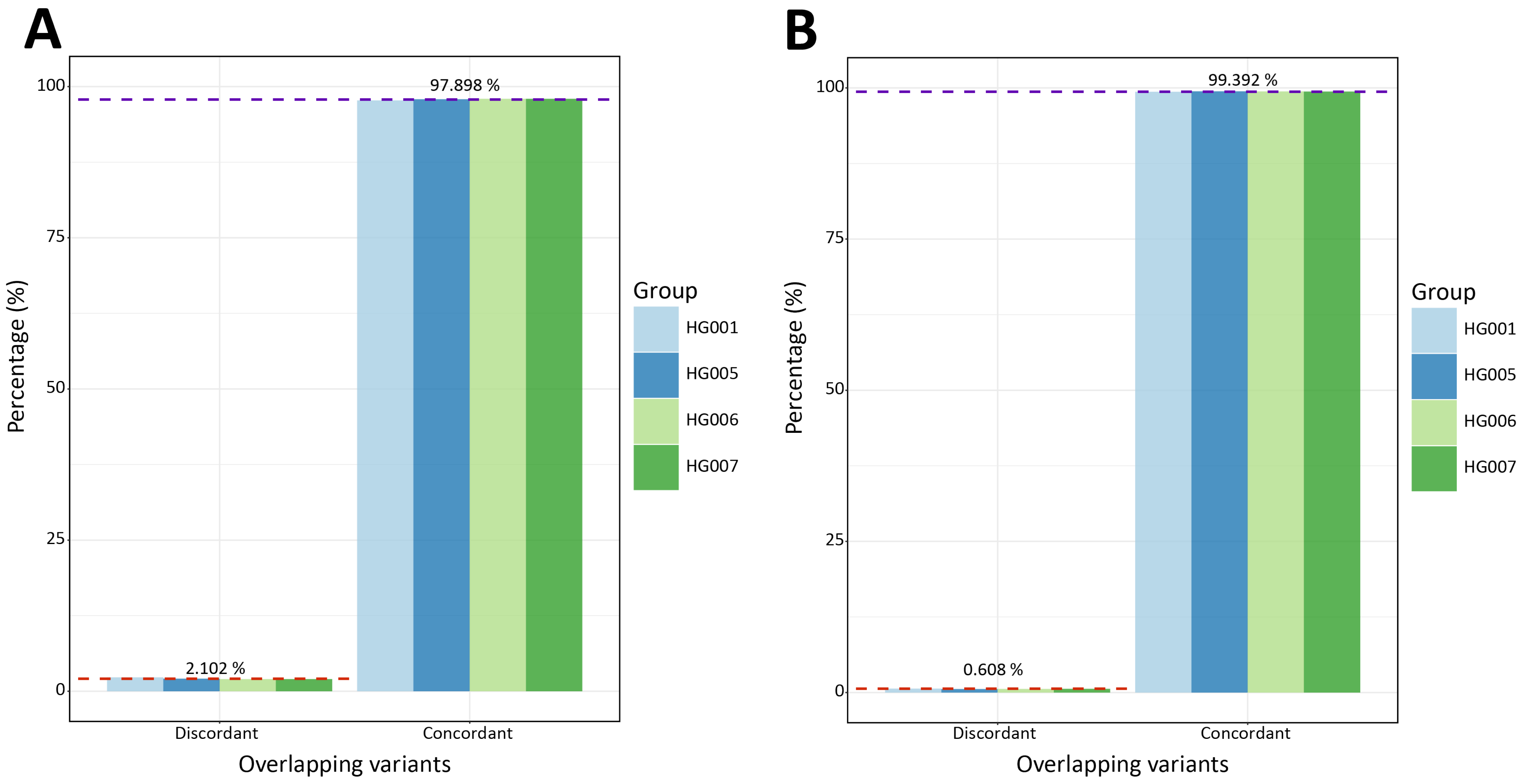

**Figure S5: Genome in a bottle (GIAB) comparison, related to Figure 2.**
(**A**) Bar plot showing the percentage of concordant (same reference and alternative allele pair) alleles and discordant alleles at matched variant sites present in the set of *n* = 48 individuals here and in benchmarked regions of *n* = 4 GIAB data sets (HG001, HG005, HG006, and HG007) [2]. Dashed red and purple lines indicate the mean percentage across all four data sets for discordance and concordance respectively. (**B**) Same as (**A**) but when only considering sites when only considering biallelic sites.

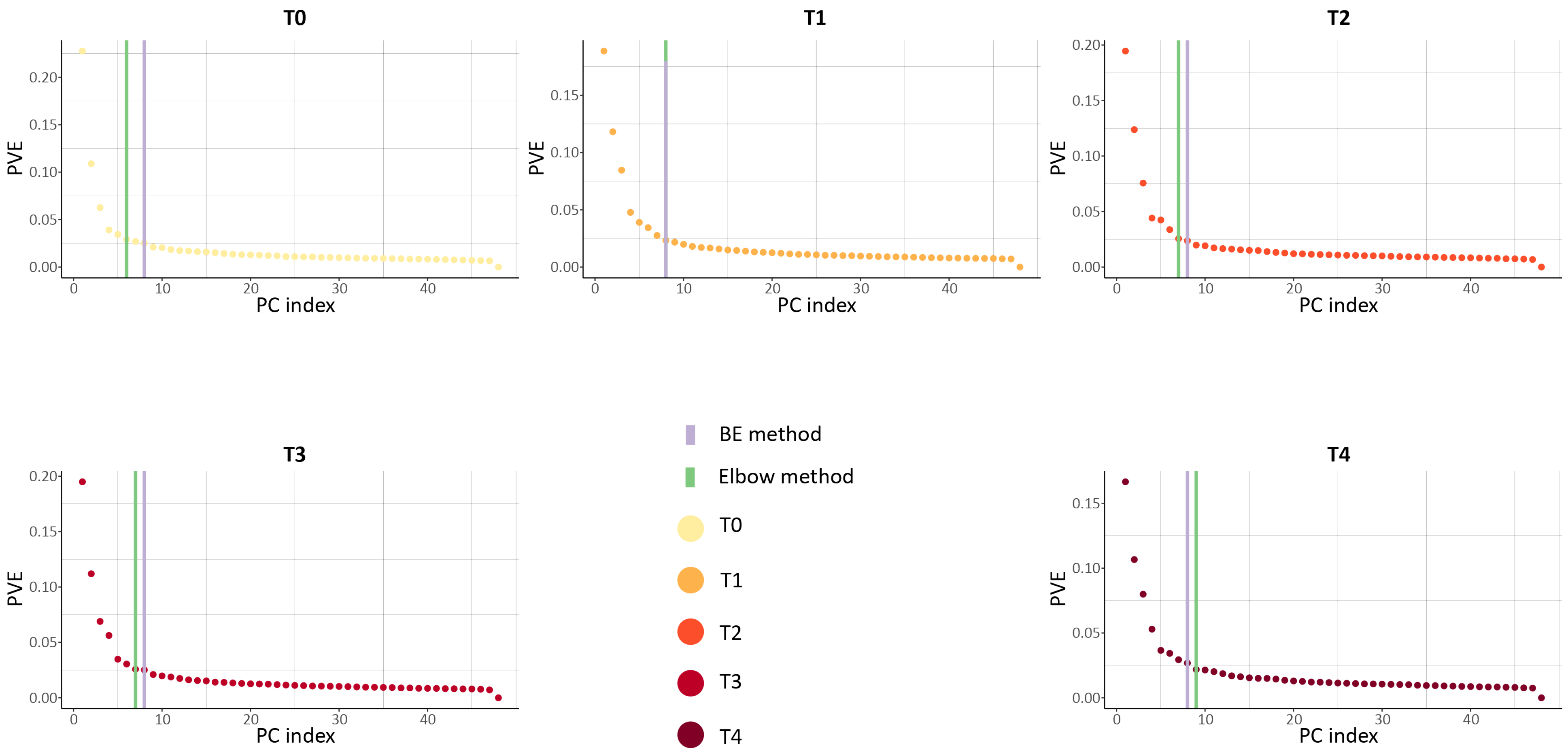

**Figure S6: Transcriptomic PCs inferred for eQTL analysis, related to Figure 4.**
Scree plots from PCA4QTL[3] for each of the five timepoints (T0–T4) showing the proportion of variation explained (PVE) by each principal component (PC) index. The green line indicates the number of PCs selected using the elbow method and the purple line indicates the number of PCs inferred using the Buja and Eyuboglu (BE) method[4].

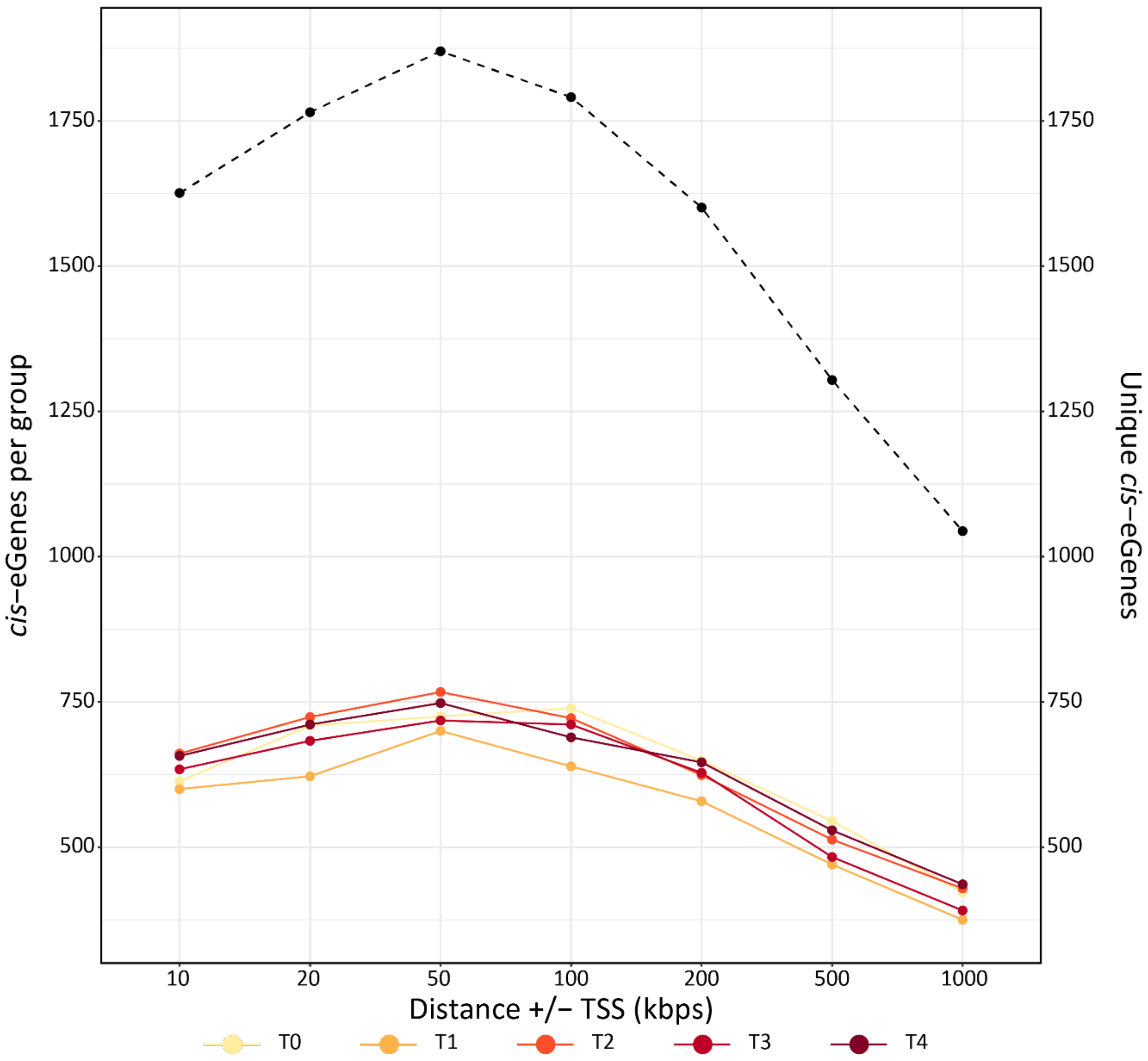

**Figure S7: Selection of optimum distance for cis-eQTL mapping, related to Figure 4.**
The number of *cis*-eGenes displayed on the first *y*-axis inferred by TensorQTL[5] for each cohort (T0–T4) using a *cis*-window size varying form 10 kb up to 1 Mb (1000 kb). The dashed line indicates the number of unique *cis*-eGenes identified across all the five groups and is illustrated by the second *y*-axis.

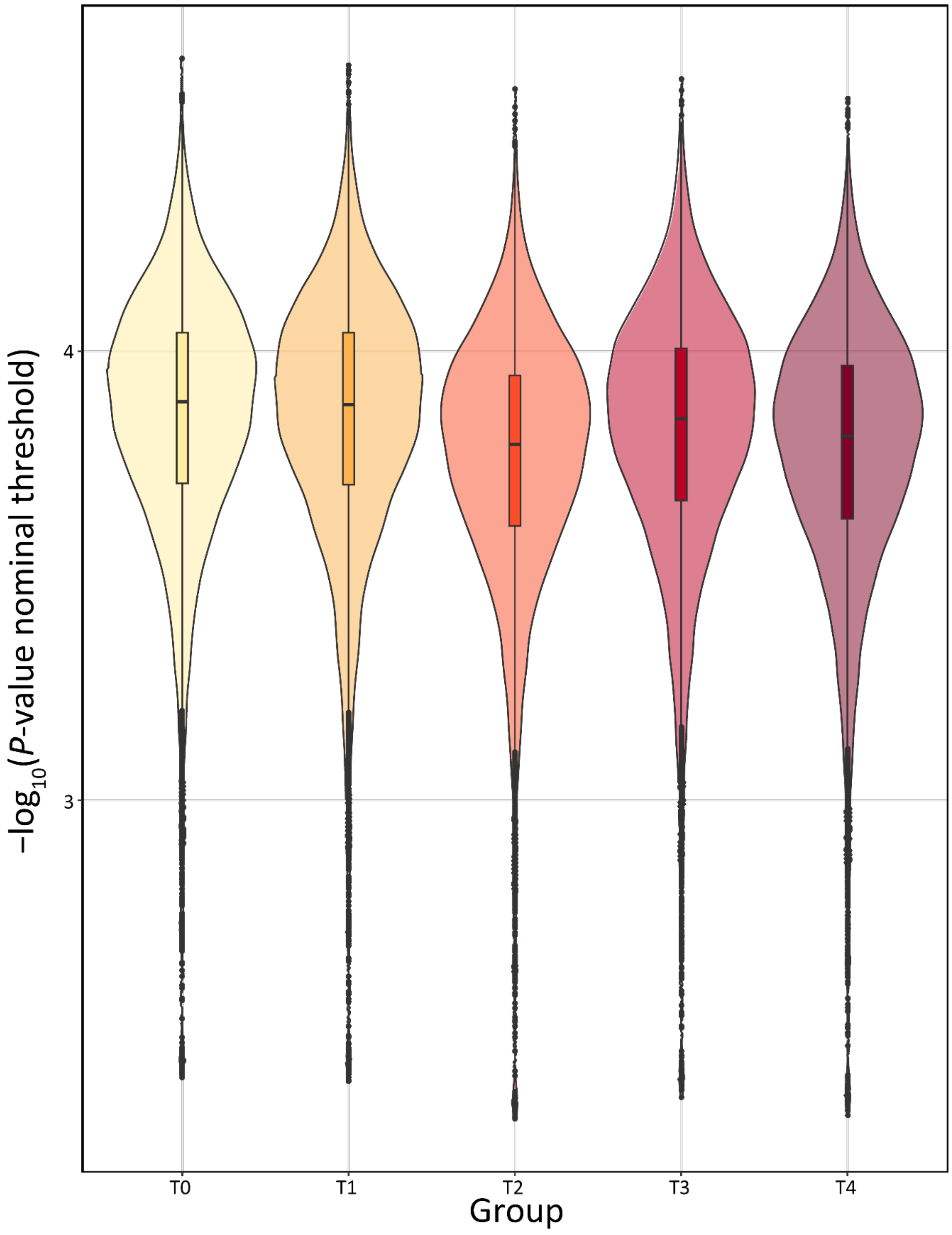

**Figure S8: Nominal P-value threshold for determining TensorQTL cis-eVariants, related to Figure 4.**

Violin plots showing the distribution of nominal *P*-value threshold per *cis*-eGene for all groups (T0–T4), respectively.

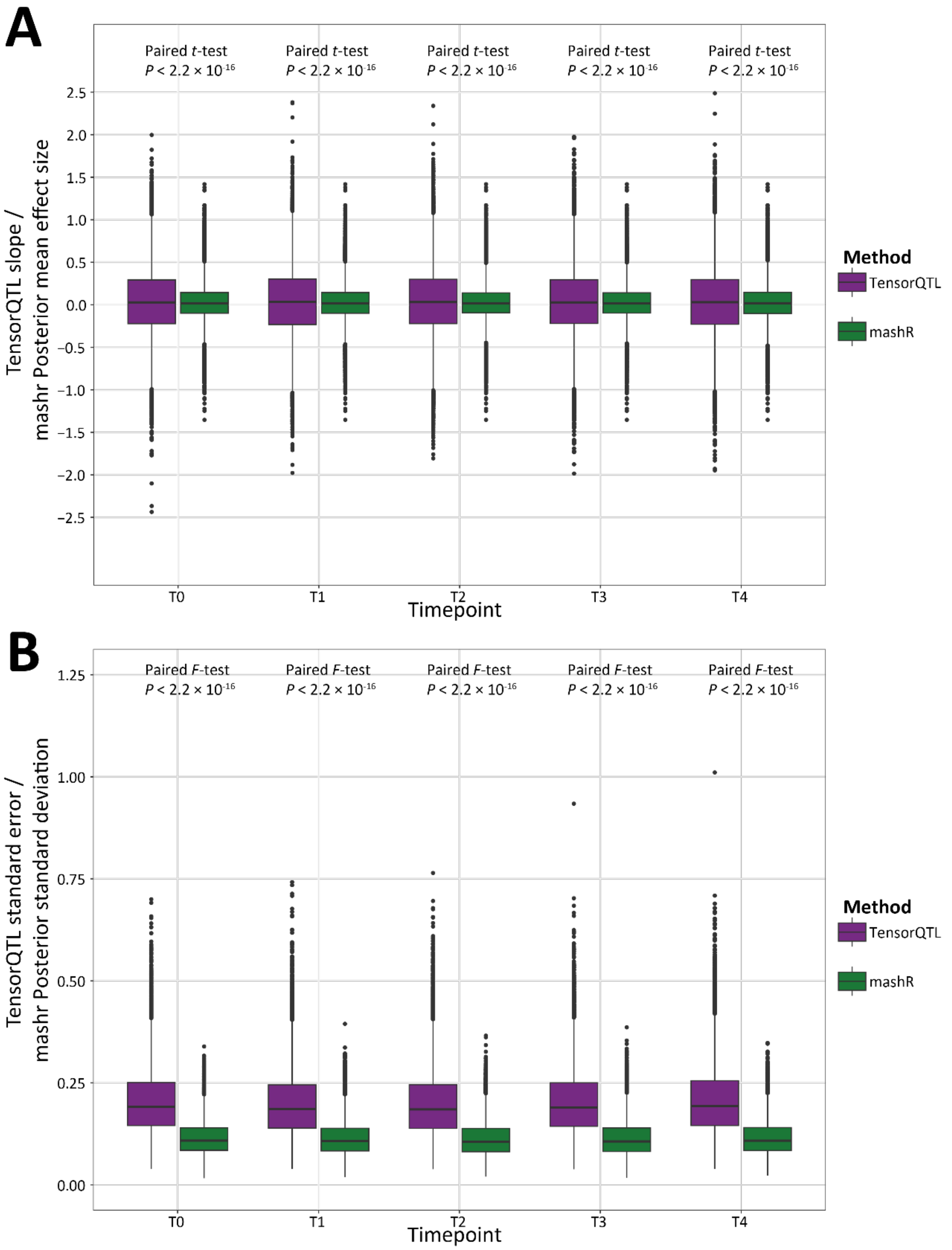

**Figure S9: Re-estimation of *cis*-eQTL effect sizes and standard error estimates using mashr, related to Figure 4.**

(**A**) Box plots showing the distribution of effect size for the most significant variant tested across all five groups and coloured based on the original TensorQTL[5] effect size (slope) and marshr[6] posterior mean effect size. *P*-values are inferred from *t*-tests comparing the mean effect size and the mean posterior mean effect size estimate in each timepoint, respectively. (**B**) Same as (**A**) but illustrating the standard error estimates for the most significant variant tested across all five groups and the mashr posterior standard deviation. *P*-values are inferred from *F*-tests comparing the standard error estimates and posterior standard deviation estimates in each group separately.

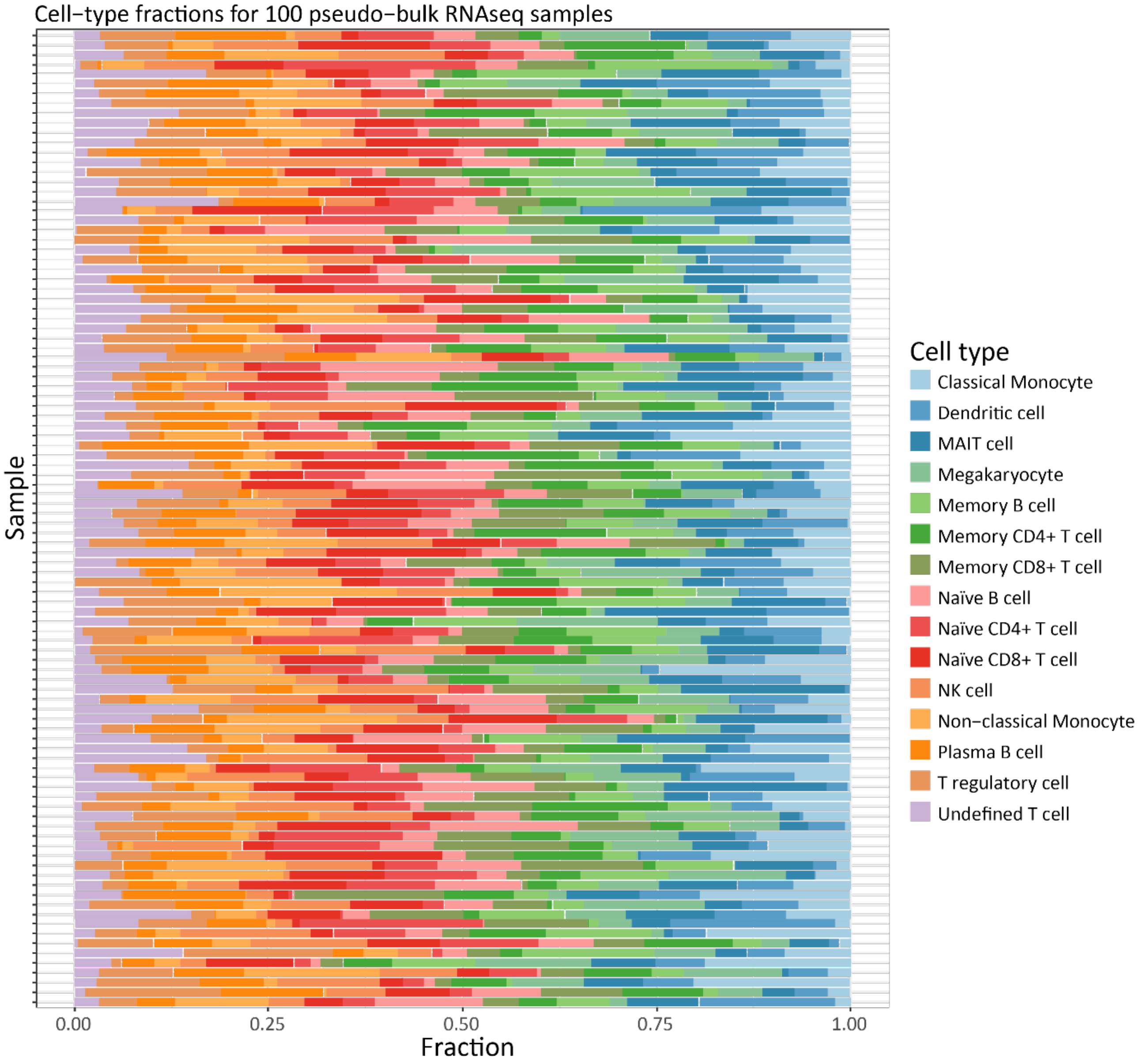

**Figure S10: Simulation of known cell type proportions using the Simbu R/Bioconductor R package based on pseudo-bulk RNA-seq data, related to Figure 5 and Supplementary Note 1.**

Bar plots of 100 pseudo-bulk RNA-seq samples on the *y*-axis with the fraction of each cell type designated and separated based on colour illustrated on the *x*-axis. Cell type proportions were generated using the Simbu R/Bioconductor R package [7] based on the scRNA-seq data used in this study.

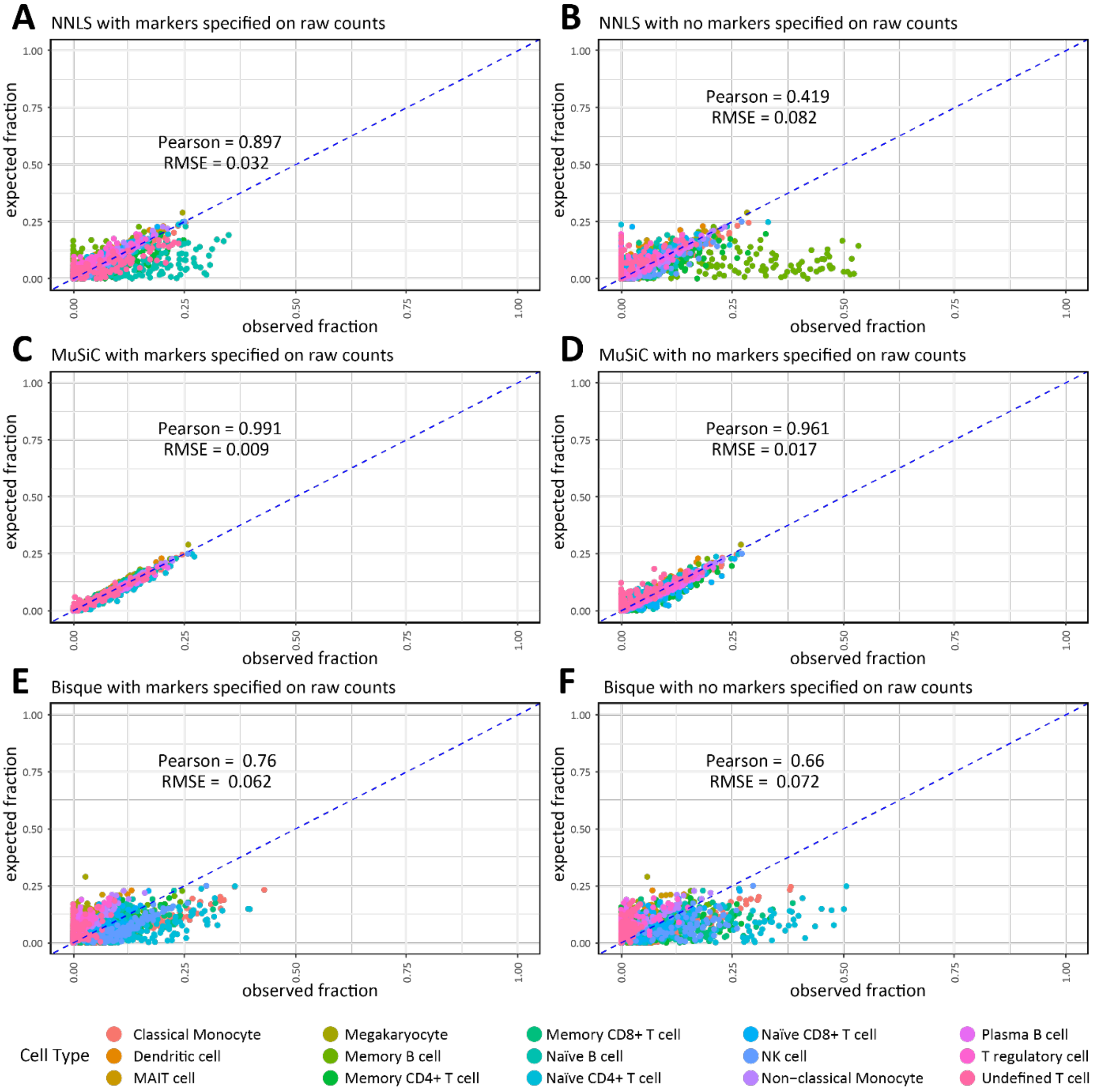

**Figure S11: Deconvolution performance of NNLS, MuSiC and Bisque with and without marker genes, related to Figure 5 and Supplementary Note 1.**

Scatter plot showing the expected fraction of cell types on the *y*-axis versus the observed fraction of cell types following deconvolution on the *x*-axis using (**A**) non-negative least squares (NNLS) with marker genes specified, and (**B**) without marker genes specified. (**C**) MuSiC with marker genes specified and (**D**) without marker genes specified. (**E**) Bisque with marker genes specified, and (**F**) without marker genes specified. Pearson, Pearson correlation for all 15 cell types; RMSE = average root mean square error for all 15 cell types.

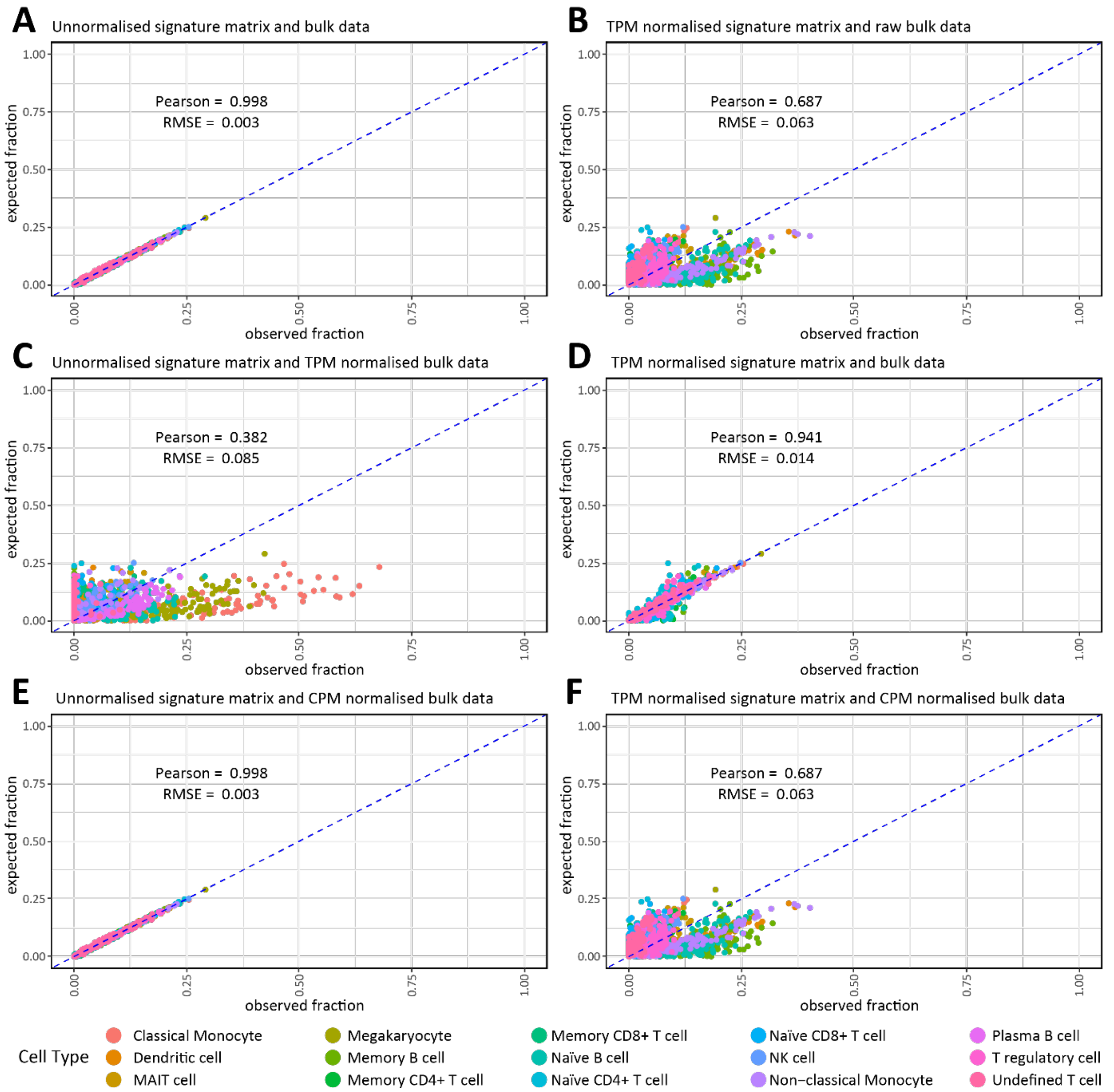

**Figure S12: Deconvolution performance of CIBERSORT with various normalisation strategies applied to the genes in the signature matrix and pseudo-bulk RNA-seq data, related to Figure 5 and Supplementary Note 1.**

Scatter plot showing the expected fraction of cell types on the *y*-axis versus the observed fraction of cell types following deconvolution on the *x*-axis using (**A**) CIBERSORT with raw unnormalised signature matrix counts and pseudo-bulk data. (**B**) CIBERSORT with transcripts per million (TPM) normalised signature matrix and unnormalised pseudo-bulk data matrix counts and pseudo-bulk data. (**C**) CIBERSORT with raw unnormalized signature matrix and TPM-normalised pseudo-bulk data. (**D**) CIBERSORT with TPM-normalised signature matrix and TPM-normalised pseudo-bulk data. (**E**) CIBERSORT with raw unnormalized signature matrix and counts per million (CPM) normalised pseudo-bulk data. (**F**) CIBERSORT with TPM-normalised signature matrix and CPM-normalised pseudo-bulk data. Pearson, Pearson correlation for all 15 cell types; RMSE, average root mean square error for all 15 cell types.

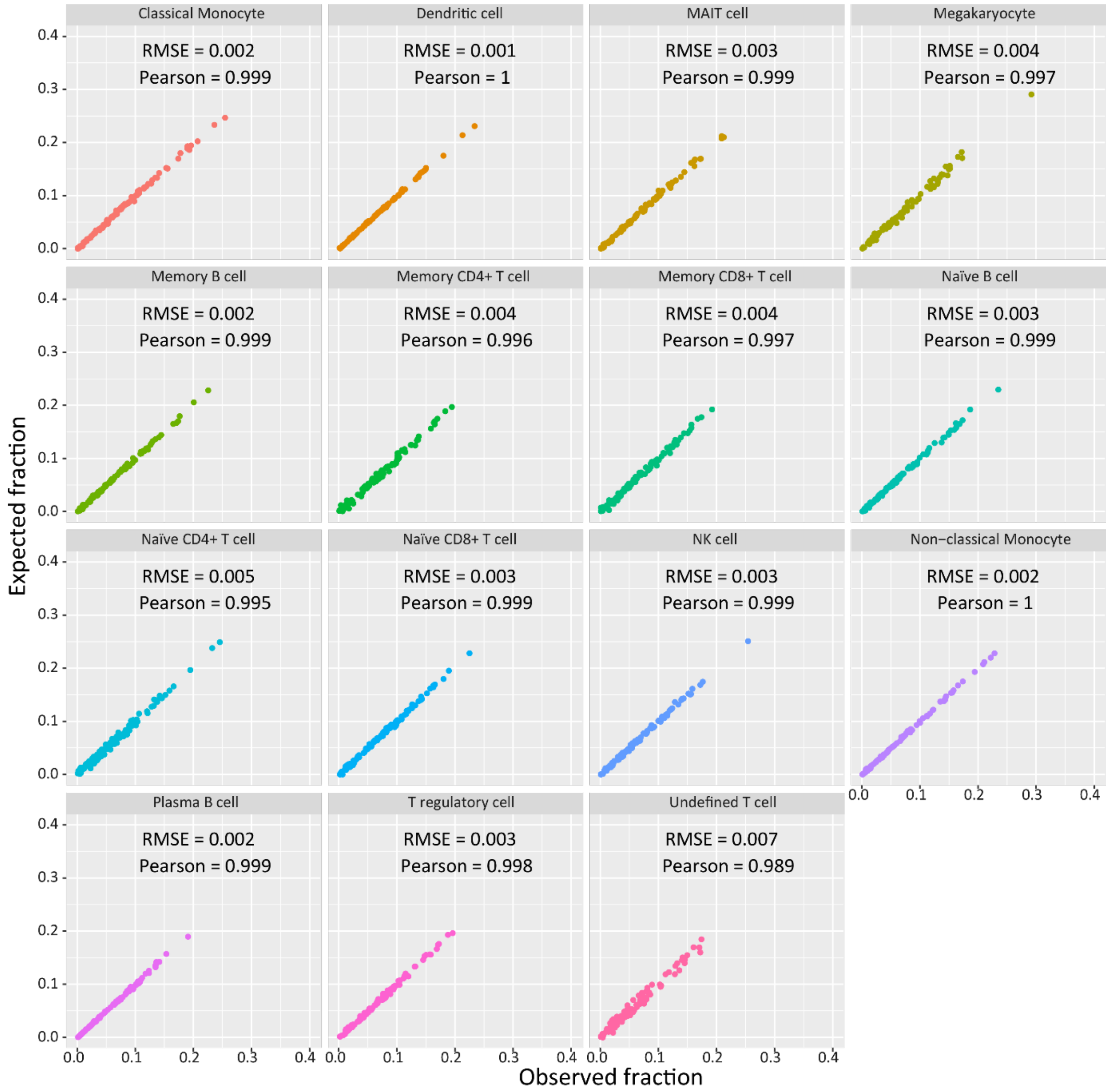

**Figure S13: Deconvolution performance of each cell type using CIBERSORT with raw unnormalized signature matrix and CPM-normalised pseudo-bulk RNA-seq data, related to Figure 5 and Supplementary Note 1.**

Deconvolution performance of CIBERSORT with raw unnormalized signature matrix and counts per million (CPM) normalised pseudo-bulk data. Pearson, Pearson correlation for all 15 cell types; RMSE, root mean square error.

### Supplementary Notes

#### Supplementary Note 1: Benchmarking deconvolution algorithms, related to Figure 5, Figure S10-13, Table S28-29.

Prior to deconvolving the PB RNA-seq data, we performed a benchmarking assessment of four deconvolution algorithms: Non-negative least squares (NNLS), MuSic, Bisque, and CIBERSORT with various parameter modifications for a total of 12 different analyses (See Methods). We then assessed deconvolution performance using the average Pearson correlation ($\rho$) and average root mean square error (RMSE) estimates across all 15 cell types. To do this, we first generated pseudo-bulk RNA-seq data for $n$ = 100 individuals derived from the scRNA-seq data with known cell type proportions using the SimBu R/Bioconductor package (**Table S28**, **Figure S11**). **Figure S12** details the results of the benchmarking assessment for MuSic, NNLS, and Bisque with, and without specifying marker genes from the signature matrix. **Figure S13** illustrates the deconvolution results for the pseudo-bulk RNA-seq data using CIBERSORT with raw or TPM-normalised scRNA-seq data and raw, CPM-, or TPM-normalised bulk data. The least well performing algorithm was CIBERSORT when specifying the raw signature matrix and the TPM normalised pseudo-bulk RNA-seq data ($\rho$ = 0.382; RMSE = 0.085). The best performing algorithm was CIBERSORT when specifying the raw signature matrix and the CPM-normalised pseudo-bulk RNA-seq data ($\rho$ = 0.998, RMSE = 0.003). We observed that CPM-normalising the pseudo-bulk RNA-seq data slightly improved the deconvolution performance but overall, had minimal impact on the performance metrics (**Figure S13**; **Table S29**). MuSiC performed well with, and without specifying marker genes ($\rho_{\text{markers(+)}}$ = 0.991, $\rho_{\text{markers(−)}}$ = 0.961; $\text{RMSE}_{\text{markers(+)}}$ = 0.009, $\text{RMSE}_{\text{markers(−)}}$ = 0.017); however, Bisque ($\rho_{\text{markers(+)}}$ = 0.76, $\rho_{\text{markers(−)}}$ = 0.66; $\text{RMSE}_{\text{markers(+)}}$ = 0.062, $\text{RMSE}_{\text{markers(−)}}$ = 0.072) and NNLS ($\rho_{\text{markers(+)}}$ = 0.897, $\rho_{\text{markers(−)}}$ = 0.419; $\text{RMSE}_{\text{markers(+)}}$ = 0.032, $\text{RMSE}_{\text{markers(−)}}$ = 0.082) were consistently poor (**Figure S12)**. For CIBERSORT with the CPM-normalised pseudo-bulk RNA-seq data, separating out the results from the deconvolution by cell type highlighted that CIBERSORT was efficient in inferring all the cell types present in the pseudo-bulk RNA-seq data ($\rho$ = 0.989–1.000; RMSE = 0.001–0.007 across all cell types, respectively) (**Figure S14**). The complete results from the deconvolution benchmarking assessment are provided in **Table S29.**
